# OpenExo: An Open-Source Modular Exoskeleton to Augment Human Function

**DOI:** 10.1101/2024.10.02.616295

**Authors:** Jack R. Williams, Chance F. Cuddeback, Shanpu Fang, Daniel Colley, Noah Enlow, Payton Cox, Paul Pridham, Zachary F. Lerner

## Abstract

While the field of wearable robotic exoskeletons is rapidly expanding, there are several barriers to entry that discourage many from pursuing research in this area, ultimately hindering growth. Chief among these are the lengthy and costly development time to get an exoskeleton from conception to implementation and the necessity for a broad set of expertise. Additionally, many exoskeletons are designed for a specific utility (e.g., ankle plantarflexor assistance) and are confined to the laboratory environment, limiting the flexibility of the designed system to adapt to answer new questions and explore new domains. To address these barriers, we present OpenExo, an open-source modular untethered exoskeleton framework that provides access to all aspects of the design process including (1) software, (2) electronics, (3) hardware, and (4) control schemes. To demonstrate the utility of this exoskeleton framework, we performed benchtop and experimental validation testing with the system across multiple configurations including (1) hip-only incline assistance, (2) ankle-only indoor and outdoor assistance, (3) hip-and-ankle load-carriage assistance, and (4) elbow-only weightlifting assistance. All aspects of the software architecture, electrical components, hip and Bowden-cable transmission designs, and control schemes are freely available for other researchers to access, use, and modify when looking to address research questions in the field of wearable exoskeletons. Our hope is that OpenExo will rapidly accelerate the development and testing of new exoskeleton designs and control schemes, while simultaneously encouraging others, including those who would have been turned away from entering the field, to explore new and unique research questions.

**One-Sentence Summary:** We present an open-source, open-hardware exoskeleton to aid in accelerating the growth of robotic exoskeleton research.

## INTRODUCTION

Exoskeletons hold immense potential as mobility enhancing and rehabilitative tools capable of transforming how, when, and where individuals engage in activity. While the field is still relatively young, it has seen a significant increase in interest within the past fifteen years. Termed “The Exoskeleton Expansion” by Sawicki and Colleagues (*1*), this increased interest has led to the development of tools with a wide range of applications including rehabilitative devices to aid those with movement deficits (e.g., cerebral palsy (*2*, *3*), stroke (*4–7*), elderly (*8–10*)), assistive devices to enhance mobility across varied activities and terrains (*11–13*), military devices to amplify soldiers’ ability in the field (*14*, *15*), and ergonomic devices to increase productivity and/or minimize injury in physically demanding workplace environments (*16–18*).

These previous devices can best be categorized using two classification schemes: (1) active or passive and (2) tethered or untethered (*1*). Active devices, those that generate assistance via powered actuators (e.g., motors), can supply higher torque magnitudes and have tunable timing that facilitate adaptive device usage but are limited by considerable added mass (vs. lighter, less adaptable passive devices). Many have overcome this limitation through tethered systems which have offboard actuators and electronics allowing researchers to ignore the tradeoff between power and mass, potentially leading to benefits beyond what is possible in self-contained systems (*19*). While valuable, these devices are restricted to the laboratory, limiting the ability to translate findings to real-world and clinical settings. Untethered (autonomous) exoskeletons have the benefit of functioning in non-laboratory settings such as at home, in the clinic, or outdoors but are limited by restrictions to torque and power due to added mass considerations. Ultimately, exoskeletons will be of most value to society when fully realized as self-contained, portable, devices; thus, the continued development of these untethered systems and their control schemes is an important future direction of the field. Another important, but often unrecognized, sub-classification of exoskeletons are upper and lower extremity devices. To date, these two types of exoskeletons have largely remained independent, with upper extremity devices focused on the shoulders (*16*, *20*) and back (*14*, *17*, *21*), and lower extremity devices mainly focused on the sagittal plane of the hips (*5*, *11*, *22*), knees (*23–25*), and ankles (*26–29*).

While there has been a large increase in activity within the field of wearable robotic devices as part of this “Exoskeleton Expansion”, the field itself is still in its relative infancy. One reason for this continued infancy is the field’s considerable barrier to entry. This barrier is characterized by a lengthy and costly developmental process that typically involves designing (1) software, (2) electronics, (3) hardware, and (4) control schemes. Each of these stages are independently challenging and often interconnected, leading to a multi-step, iterative process that requires a substantial investment of time and resources before novel research can be performed. An additional barrier is that the field is relatively inaccessible to non-experts. That is, to develop an exoskeleton device, one needs expertise in a broad set of domains, including mechanical and electrical engineering, robotics and controls, and biomechanics, which can be prohibitive for those lacking experience in one of these disciplines (e.g., biomechanist, physical therapist, computer scientist). In addition to having a high barrier to entry, the field is also limited by the use of highly specialized, organizationally-embedded systems. This limitation manifests in several different ways. First, the creation of highly specialized devices (e.g., a device solely focused on ankle plantarflexion) leads to inefficiencies when a researcher wants to explore a new domain (e.g., hip or elbow joint) as much of the software and hardware utilized in one device will have to be adapted to interface with a new device. Second, there is a large variety of organization-specific software and hardware that groups use to conduct their research and contribute to the field. The problem with these closed-source systems is that it results in increasingly isolated and specialized systems that make research increasingly harder to replicate, further contributing to the reproducibility crisis faced by much of the research community (*30*). Finally, these highly specialized, closed-loop, devices typically lead to small sample sizes during experimental testing which increases the likelihood of irreproducible work. One solution to these challenges would be the establishment of a fully open-source exoskeleton system. Similar devices in other fields, such as in powered prosthetics (*31*) and in quadrotor flight (*32*), have proven to be valuable tools to new and established researchers. Some have tried to develop open-source exoskeleton devices before, such as OpenBionics’ exoskeleton glove (*33*), FlexSEA (*34*), and the pediatric-focused ALICE (*35*), but these devices are highly specialized and thus relatively inflexible and therefore less likely to be widely adopted. All these challenges and limitations could be addressed by developing a fully open-source system that is accessible to those without complete expertise in the field while simultaneously being flexible enough to facilitate new device designs with minimal changes needed to the system’s architecture.

To address this, we have created OpenExo, a modular open-source exoskeleton system (software, hardware, electronics, and controls) for researchers and curious global citizens to access, utilize and expand as they see fit. Here we present the design of the software, electrical architecture, and example hardware interfaces of this open-source system, characterize performance under different configurations, and demonstrate modularity by assessing its functionality under a variety of assistive configurations (hip-only, ankle-only, hip-and-ankle, and elbow). Our hope is that our community and those who wish to enter the field will embrace OpenExo, utilize it to interface with novel joint assembly end-effectors, and bring us closer to making exoskeletons a reality of our everyday life.

## RESULTS

### Open-Source Tool

All resources related to OpenExo (**Fig. 1A**) can be found at theopenexo.org. This includes: (1) a software package capable of use with or without modification to support the rapid testing of new hardware and control approaches without the need for extensive development of supporting code (**Fig. 1B**), (2) a detailed electrical architecture (**Fig. 1C**), including a printed circuit-board (PCB) designed to interface with the software package while performing bilateral operation of up to two joints at a time (e.g., hip-only, hip-and-ankle, hip-and-elbow), and (3) an open-source hip design (**Fig 1D**) and Bowden-cable transmission system (**Supplemental Figure 1**). Importantly, these resources are supported with extensive documentation to help facilitate usage and modification for experts and novices alike. This includes a guide to the structure and function of the software and how to perform key modifications, such as adding new joints, controllers, sensors, and motors, as well as a guide to introduce novices to the programming language utilized throughout the code (C++). This also includes information on running and utilizing a companion python application (included as part of the software) capable of facilitating device operation while giving users the ability to modify control parameters (e.g., applied torque magnitude) and monitor/store data in real-time. We have also detailed the structure and function of the PCB, included information on how and where to get it externally manufactured, and provided the files needed to enable researchers to modify it to support their own needs. Finally, we have provided full supporting documentation for the construction of a low-cost direct-drive hip device and a Bowden-cable transmission system. This includes the part files, a complete bill of material with a cost breakdown and where to purchase materials, a step-by-step guide to manufacturing and constructing all components involved with these devices (machining and assembly of parts, creation of wires and cables, how they all fit together), and directions to 3^rd^ party resources for those who lack the means to manufacture the hardware in-house.

**Fig. 1.**
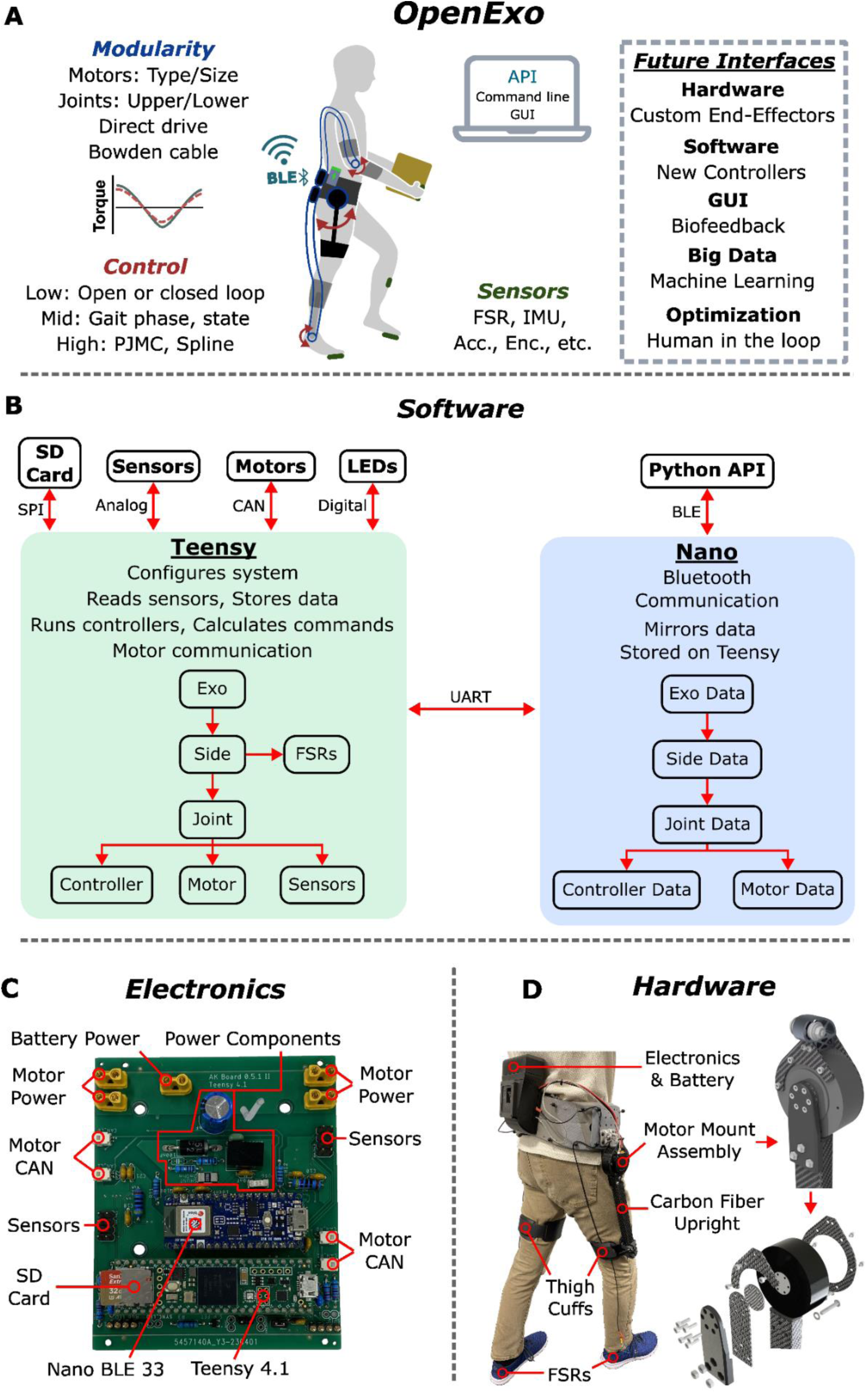
Open-source software structure, electrical architecture, and hip hardware. **(A)** Overview of OpenExo, emphasizing its modular features including: swapable motors, flexible joint and actuation schemes, varied control approaches and sensors, and a companion API. Future planned interfaces include custom end-effectors, new controllers, updated GUI features, and integrated machine learning and human-in-the-loop optimization approaches. **(B)** Breakdown of the structure and function of the software package. The software was designed to have a high degree of modularity while minimizing redundancy. It operates on two microcontrollers simultaneously (Teensy 4.1 and Arduino Nano BLE 33) to facilitate both computation and Bluetooth communication while maintaining a fast operation speed. Detailed guides on the software are available to aid researchers in effectively utilizing this resource to meet their needs. In addition, a companion python Bluetooth application is also available to help users operate and control the system in real-time. **(C)** Breakdown of the printed circuit board used for device operation. It supports both microcontrollers needed for device operation and has connections to facilitate bilateral operation of up to two joints at a time. **(D)** Breakdown of the components to the direct-drive hip device. Detailed guides to manufacturing and assembling the device are available to help researchers replicate the design.

### Design Goals

There were several design goals that went into the creation of this system. This included creating a device that (1) could be utilized in any setting (e.g., indoor, outdoor, mixed terrain), (2) had a logical and easy to follow software structure that was accessible to experts and novices alike, (3) had a simple, yet robust, electrical architecture that can be utilized to support both single and multi-joint assistance while being easily revisable to meet researcher’s evolving needs, (4) had a hardware system that could support multiple configurations simultaneously, all of which could be easily swapped depending on the user’s desired task, (5) included an easy to manufacture, low-cost base hip configuration to enable novices to replicate our work and begin developing their own projects, (6) had a Bowden-cable transmission system that could be used as a starting point for others to pursue innovative joint assembly end-effector designs, and (7) embedded modularity into all aspects to facilitate rapid testing of new hardware and control schemes without the need for extensive modification of the software and electrical components. To summarize, we sought to design a system that minimized redundancy and could support multiple joints at a time without limits to what joints could be included in the system (e.g., both lower and upper extremity joints) while being capable of being used by anyone in any setting. To achieve this, we embedded principles of modularity in all components of the system including its software, electrical architecture, and hardware.

### Software

The chief source of modularity in our open-source system comes from the software architecture. Our software was developed using a combination of C++ and Arduino languages, with the principle of inheritance-based polymorphism (realized through parent-child and abstract classes) used to achieve a high-degree of modularity, reduce redundancy, and minimize dependencies. **Figure 1B** outlines the structure of the software, which was designed to have a logical and easy to follow form. The system starts broad and develops a narrower focus as it proceeds through its computations, starting with functions related to overall exoskeleton usage, accessing computations specific to each side, then getting into computations specific to joints which includes accessing the controllers, sensors, and motors assigned to those joints. Importantly the controller, sensor, and motor computations are all self-contained and accessed by instances of joints. That is, if you have a controller capable of operating on multiple joints you don’t need to repeat the controller definitions for each joint, they can be defined once and accessed by any joint that can operate said controller.

To avoid having to directly modify the software every time a change in device configuration was desired, we designed the system to pull in the desired exoskeleton configuration information from a file stored on a secure digital (SD) card that slots into a Teensy microcontroller. Through this, users can change which joints and sides of interest they would like to run as well as which controller is the default mode of operation for the specified joint. Additionally, we have developed a companion Bluetooth application programming interface (API) in python to allow users to update controller parameters and monitor/store data during operation.

### Electrical Architecture

Our focus was on developing simple and intuitive electronics to facilitate fabrication of an exoskeleton device among individuals who may not necessarily have expertise in electrical design (e.g., biomechanist, computer scientist). Previous open-source electrical architectures, such as FlexSEA (*34*), are extremely modular but also rather complex due to the usage of multiple interconnected PCBs which may limit more widespread adoption and discourage researchers from attempting revisions to meet the needs of their device. We opted for a simpler approach that encourages iteration and revisability to help facilitate longer-term adoption and thus reduce the need for repeated work that occurs with redesigns.

Our PCB consists of one board that can support up to two joints, bilaterally, at a time. We developed the board to interface with CubeMars’ AK-series motors (Nanchang, China) as they are powerful enough to provide torques for larger adult participants and during more challenging tasks such as multi-terrain walking (e.g., stairs, inclines, and declines). Additionally, there are multiple versions of these motors which all rely on the same communication scheme (Controller Area Network; CAN). Thus, by designing the electronics and software to interface with CAN communication, researchers can choose which version of the AK Series motors to work with, further enhancing modularity. For example, if a researcher wanted to utilize the device on a pediatric individual or during a low-demand task, then the AK60-6v1.1 motor could be used as it has smaller torque generating capabilities while being much lighter than other versions of the motor. If a researcher wanted to use the device on an adult participant or during a higher-demand task, they could switch to the AK80-9 or AK70-10 versions of the motor to get higher torque generation at the tradeoff of added mass. The board was also designed to take an eight-pin ribbon cable connector on each side, providing room for multiple external sensor connections (e.g., force sensitive resistors [FSRs], torque transducers, angle sensors) that can aid in device control and sensing. We have outlined the process for modifying the software to take new boards, sensors, and motor types should users wish to modify the system to perform new functions or explore new actuation approaches.

### Hardware

The software and electrical architectures were designed to be modular so that researchers could develop hardware capable of assisting any upper- or lower-extremity joint of interest. To highlight this, we developed numerous different configurations to interface with this system. This included hardware to aid the hip (**Figure 2A**), ankle (**Figure 2B**), and elbow (**Figure 2D**). All of these were designed to interface with the same waist-belt and are capable of being combined (e.g., hip-and-ankle assistance; **Figure 2C; Supplemental Figure 2**) and swapped quickly to enable researchers to rapidly explore new types of exoskeleton assistance (e.g., combined hip-and-elbow assistance) with minimal re-tooling necessary. To facilitate easy replication by non-experts, the hip hardware was designed to be simple, easy to manufacture, low cost (∼$2,000 – 2,500), and capable of operating accurately under open-loop control. Additionally, we have included the design of our belt-side Bowden-cable transmission so that others may replicate and explore new end-effector design approaches via this form of actuation (**Supplemental Figure 1**). In all cases, the hardware was designed to operate only with the electronics embedded in the waist belt to enable researchers to utilize the device in both indoor and outdoor settings.

**Fig. 2.**
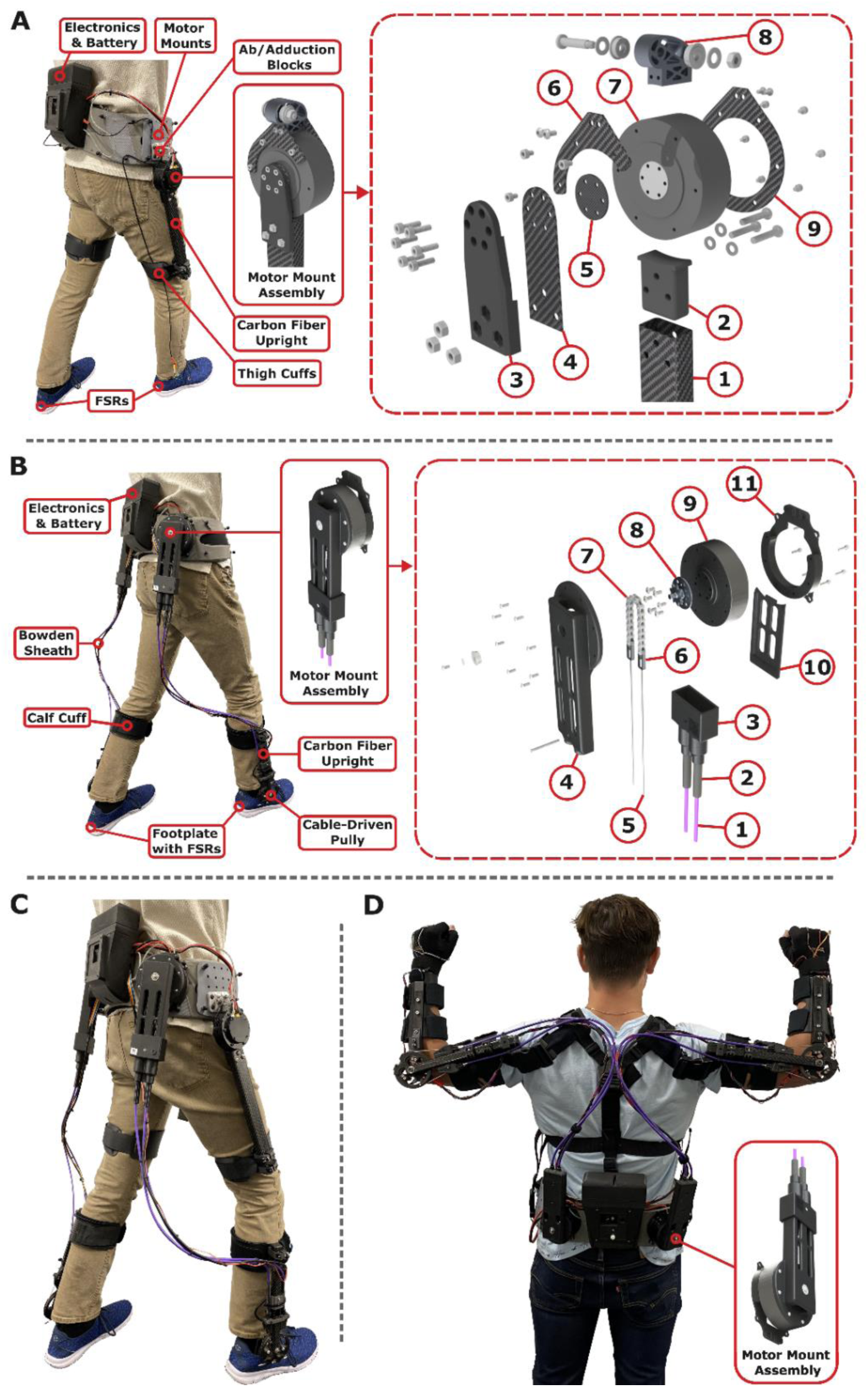
Different exoskeleton configurations operating on OpenExo’s software and electrical architecture. **(A)** Direct-drive hip exoskeleton, **(B)** Bowden cable-based ankle exoskeleton, **(C)** Combined hip-and-ankle exoskeleton, **(D)** Elbow exoskeleton with the motor mount Bowden-cable transmission highlighted to emphasize the flexibility of the belt-to-motor interface design, which is just the inverted interface utilized for the ankles.

The design of this transmission system is openly available for others to replicate and utilize (**Supplemental Figure 1**). Configurations are combined on the belt (**Supplemental Figure 2**) and connected to the PCB secured in the Electronics and Battery housing in the middle of the waist belt. A detailed breakdown of the exploded view/parts labels is outlined in the **Supplemental Material**.

### Engineering Validation

We quantified the capabilities and limitations of the system by completing benchtop and performance tests for each configuration. This included benchtop testing to characterize the responsiveness of the device to changes in torque, torque tracking tests during functional operation to understand the accuracy of the prescribed assistance, and duration testing to gain insight into the limiting factors surrounding device lifespan.

### Benchtop Testing

We characterized the responsiveness of each individual joint when subjected to a step-response at torque magnitudes realistic to those encountered while operating the devices. To achieve this, all devices needed an in-line torque transducer so that applied torque could be measured and compared to what was prescribed. This required a small change in the design of the open-loop hip hardware to facilitate this measurement (**Supplemental Figure 3A**). In all cases, the device operated with the low-level control scheme intended for use while worn by a user. That is, the hip device, despite the presence of the torque transducer, operated under open-loop control, while the ankle and elbow devices operated under closed-loop control. While subjected to an open-loop step response of 6 Nm, the hip device displayed excellent responsiveness, with a rise time of 3.0 ms and an overshoot of 0.30 % (**Figure 3A**). During closed-loop control, the ankle configuration also displayed a fast response, with a rise time of 65.0 ms and an undershoot of 1.8% when subjected to a step response of 28 Nm (**Figure 3A**); while the elbow configuration had a rise time of 35.8 ms with an overshoot of 6.4% under a 10 Nm step response (**Figure 3A**) (*36*).

**Fig. 3.**
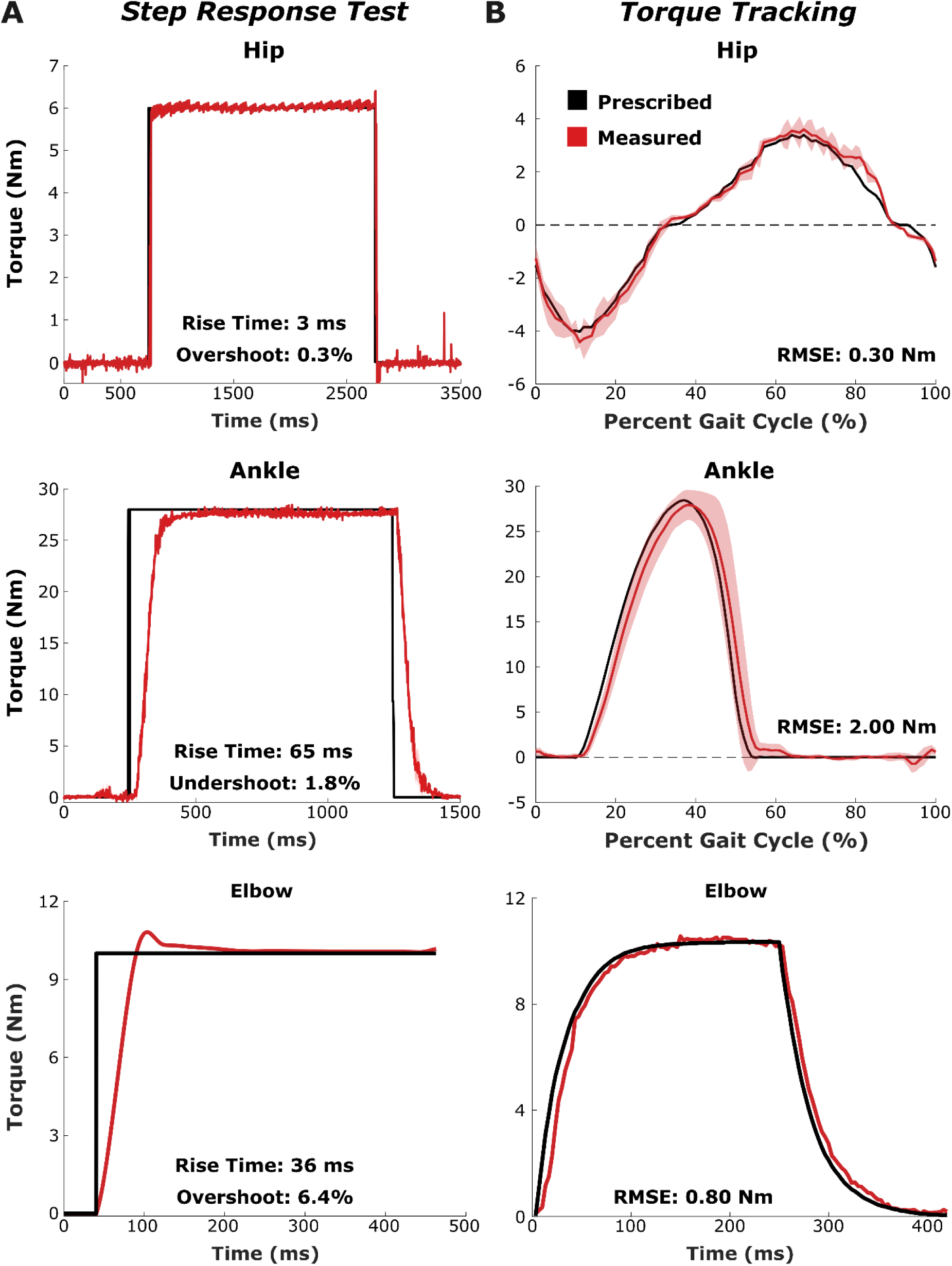
Results from the benchtop engineering validation indicate the system has a fast and accurate response regardless of configuration. (A) Step response test for the hip (top), ankle (middle), and elbow (bottom) configurations. Each configuration displayed fast rise times with minimal overshoot/undershoot. (B) Torque tracking test for the hip (top), ankle (middle), and elbow (bottom) using the same control schemes as the experimental validation. Each configuration had low tracking error (∼7% of the maximum torque magnitude).

### Torque Tracking

To characterize the accuracy of the prescribed assistance, individuals donned and operated the device during their intended function (i.e., walking for the hip and ankle configurations, weightlifting for the elbow) while the measured and prescribed torques were recorded. While in the hip configuration operating on a controller developed by Bryan et al. (*19*) and Franks et al. (*37*), the root-mean squared error (RMSE) of the device was 0.30 Nm (7.3% of the prescribed setpoint), suggesting that the control paradigm was accurate despite the lack of closed-loop, low-level control (**Figure 3B**). While configured for ankle assistance using a previously described proportional joint moment controller (PJMC) (*29*, *38*), the device maintained this same level of accuracy, despite the Bowden-cable transmission system, with a RMSE of 2.00 Nm (7.1% of the setpoint; **Figure 3B**). Finally, while assisting the user during a repetitive lifting task via a novel controller (*36*), the elbow configuration displayed a similar accuracy as the hip and the ankle with a RMSE of 0.84 Nm (7.0% of the setpoint; **Figure 3B**).

### Duration Testing

To fully characterize the utility of a device, it is important to understand how long it can operate and its limiting factors. Thus, a duration test was performed with a user operating the device in each configuration while using a 22.2v, 1800 mAh Li-Ion battery (HRB). For the hip, ankle, and hip-and-ankle configurations this involved walking on a treadmill at 1.25 ms^-1^ while each joint operated at the user’s maximum comfortable torques. As the device operated in these conditions the total battery voltage and the motor temperature on both sides were recorded at one-minute intervals. The test was halted once the motors reached their temperature limit (100 °C) or once the battery voltage reached the manufacturer’s recommended stopping voltage (3.7 V/cell = 22.2 V). The open-loop hip configuration (using AK60-6v1.1 motors) was able to operate for 35 continuous minutes at peak assistance. During this time the average motor temperature increased without reaching a steady state while the battery voltage decreased steadily (**Figure 4**). At the end of the walk the motors had reached a temperature of 76.7 °C while the battery had reached the manufacturer’s recommended limit of 22.2 V. When walking with peak plantarflexion assistance, the ankle configuration (using AK80-9 motors) operated for 25 continuous minutes. Similar to the hip, the motor temperature continuously increased, but failed to level off, reaching a maximum temperature of 56.8 °C while the battery decreased until reaching the manufacturer’s recommended limit (**Figure 4**). When configured to provide simultaneous assistance to the hip and ankle joints, with maximum assistance at each joint, the device was able to operate continuously for 15 minutes. Like the hip-only and ankle-only configurations, the motor temperatures failed to level off, reaching a temperature of 56.7 °C at the hips and 40.8 °C at the ankles, while the battery reached the manufacturer’s limit (**Figure 4**). It should be noted that, while the manufacturer recommends a limit of 3.7 V/cell, this is with long term health of the battery in mind and likely represents a conservative estimate of battery life for a given session. For single operation, it is likely that the battery could be taken down to ∼3.2 V/cell (total: 19.2 V) before reaching its true limit, and thus these duration tests were likely ended far earlier than necessary. As a result, the battery voltage is presented as a % capacity and is terminated at ∼50% of the capacity (**Figure 4**).

**Fig. 4.**
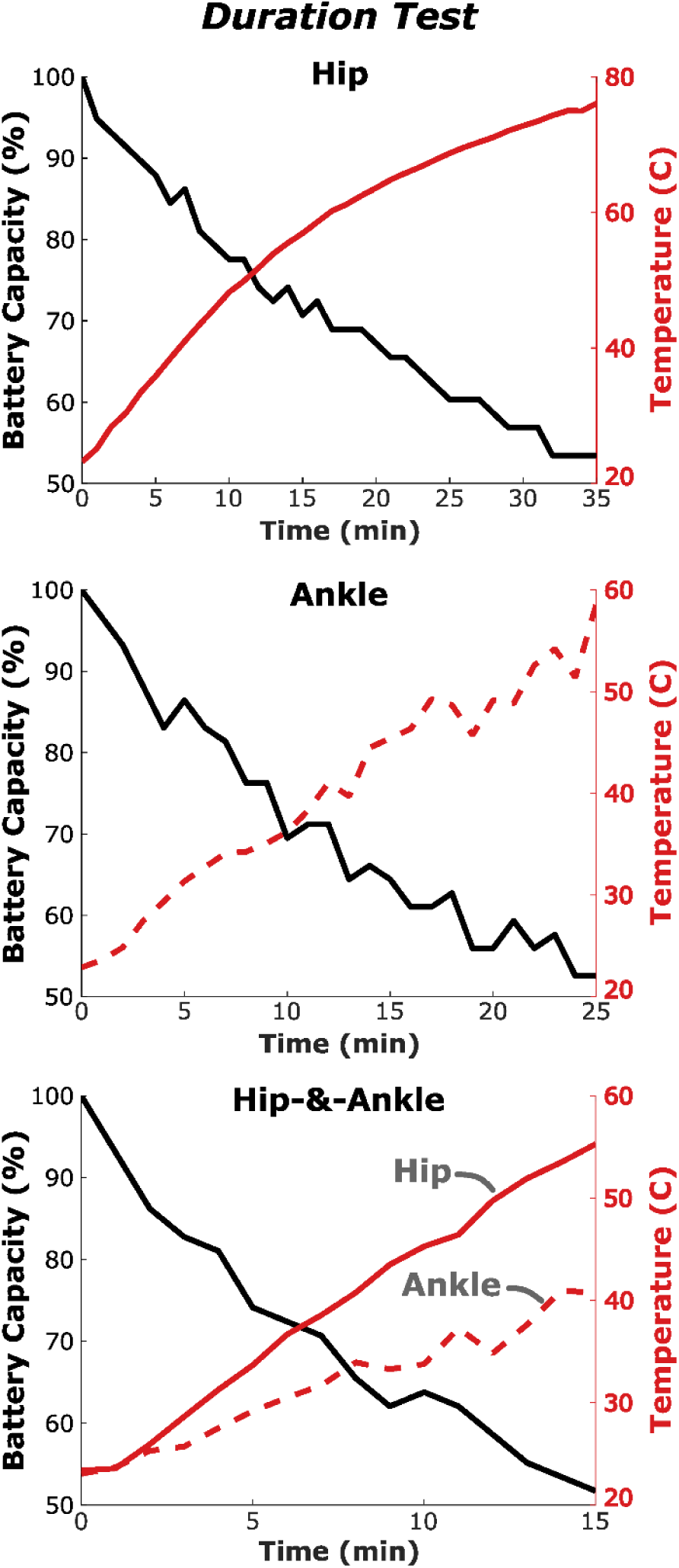
Duration testing of the OpenExo under its lower-extremity walking configurations indicates operation lifespan is primarily limited by battery life. Battery capacity (black) and motor temperature (red) over the course of a duration test for the hip (top), ankle (middle), and combined hip-and-ankle (bottom) configurations. The tests were ended when the battery voltage reached the manufacturer’s recommended limit of 3.7 V/cell (22.2V total), which represents 50% of the battery’s actual estimated capacity (3.2 V/cell = 19.2 V total) or when the motor temperature reached their shutoff temperature (100 °C). All configurations were operated at their maximum comfortable torque setpoints while walking on a treadmill at 1.25 ms^-1^. The hip lasted 35 minutes, the ankle 25 minutes, and the combined hip-and-ankle 15 minutes. In each case, the battery was the limiting factor, suggesting this could be a primary target to improve device operation lifespan moving forward.

The elbow configuration proved challenging to estimate an operation time. The device was designed for a user to wear in a physically demanding workplace environment where they would need assistance for some tasks and not others. Thus, the control scheme was designed to give the user control of when the device was engaged so that the device could be worn and functional for all tasks encountered in the day-to-day workplace environment. This presents a challenge for estimating the device’s single-use lifespan as it is hard to operate in a consistent cyclical manner (which would likely never be encountered in the workplace). Thus, to estimate the overall runtime of the device in this configuration, we had a user simulate a plausible workplace task while operating the device as they would in that environment. Specifically, we had the individual transfer 10 kg boxes between two shelves while the device provided 12 Nm of assistance per arm during each carry, with an estimated 30% of device downtime (where it did not provide assistance) during the walk back to the original shelf without a box to carry. This was performed for a ten-minute period, where the initial voltage was 25.18 V and the final voltage was 24.78 V, representing a 7.6% decrease. Extrapolating this rate of power consumption, it would take ∼2.2 hours of operation (using AK60-6v1.1 motors) until the battery reached the manufacturer’s recommended limit (representing 50% of its potential capacity), suggesting that this system would likely be functional over the course of an entire day’s worth of usage (*36*). Motor temperature was not recorded during this task as the battery voltage was the limiting feature of every other configuration and recording the temperature during this task would disrupt the task itself, influencing the results.

### Experimental Validation

To demonstrate the versatility and utility of our open-source system, we recruited healthy adults (n = 7, **Supplemental Table 1**) to complete activities while the device operated in its different configurations. We selected activities for each configuration that we thought would elicit the most benefits from device usage. This included incline walking while configured for hip assistance (n = 2), as the hips contribute more to positive power generation during incline walking (*39*). This also included outdoor and indoor walking while configured for ankle assistance (n = 1), as the ankles contribute the most positive power during level walking (*39*). We sought a more challenging, high-demand, task for multi-joint assistance and thus selected load carriage during level walking for the combined hip-and-ankle configuration (n = 2). Finally, we had users lift weights to fatigue (n = 2) while the device was configured for elbow assistance, as the intended function of this configuration was to assist in lifting tasks.

### Hip – Incline Walking

While configured for hip assistance, we had two individuals (**Supplemental Table 1**; P1 & P2) perform incline treadmill walking at 7.5^◦^ while outfitted with a portable, indirect calorimetry metabolic unit (K5, COSMED). Both individuals walked for eight minutes for each condition from which the metabolic-power for the last two minutes was calculated and then normalized by body mass and walking speed to determine the steady-state metabolic cost of transport (*29*, *40*). Both participants displayed reductions in the metabolic cost of transport while using the hip exoskeleton, compared to walking without the exoskeleton, during incline walking (**Figure 5A**). P1 had an 8% reduction in metabolic cost (Shod: 6.72 Jkg^-1^m^-1^; Exo: 6.17 Jkg^-1^m^-1^), compared to the shod condition, while P2 had a 14% reduction (Shod: 7.35 Jkg^-1^m^-1^; Exo: 6.29 Jkg^-1^m^-1^).

**Fig. 5.**
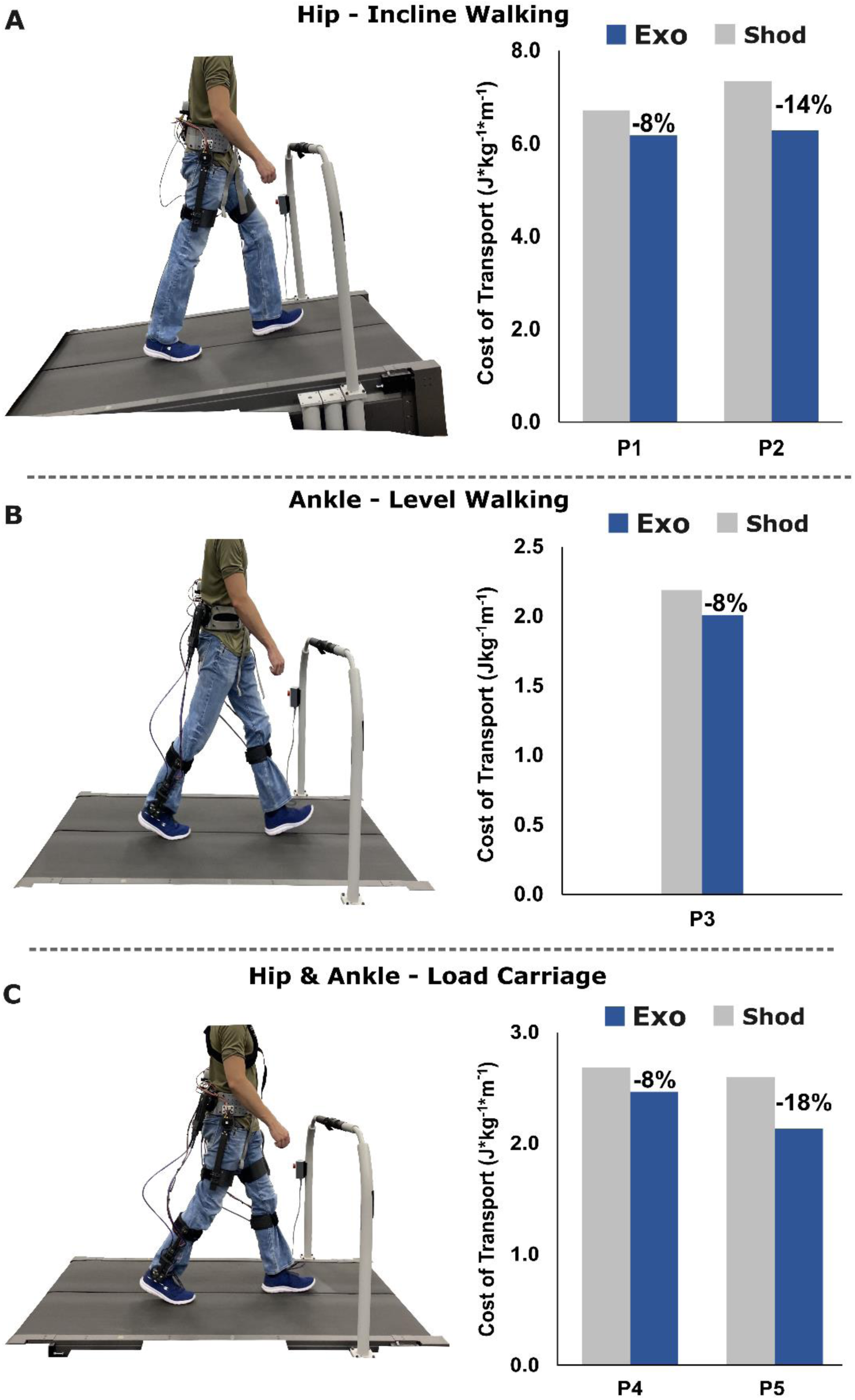
Metabolic cost of transport for each of the experimental treadmill tasks was lower while receiving assistance when compared to walking without the device. (A) Metabolic cost of transport with (blue) and without (grey) hip assistance during 7.5^◦^ incline walking at a self-selected speed. (B) Metabolic cost of transport with (blue) and without (grey) ankle assistance during level walking. (C) Metabolic cost of transport with (blue) and without (grey) combined hip-and-ankle assistance while walking with a weighted vest. All three images on the left are representative images of a user walking with the specified configuration during the task of interest.

### Ankle – Outdoor & Indoor Testing

One individual (**Supplemental Table 1**; P3), with prior ankle exoskeleton usage experience, completed ankle-assisted walking on outdoor terrain. This involved walking with, and without, the ankle exoskeleton about a relatively flat 1650 m loop at a local park (**Figure 6A&B**). The time to complete the loop and the number of steps taken with the right leg were recorded and used to calculate the average speed and stride length of the user during each condition. While walking outdoors in the exoskeleton (vs. without the exoskeleton), the user completed the loop faster and with fewer steps, resulting in a longer average stride length and a faster average walking speed (**Figure 6C**), suggesting improved mobility while using the device in this configuration.

**Fig. 6.**
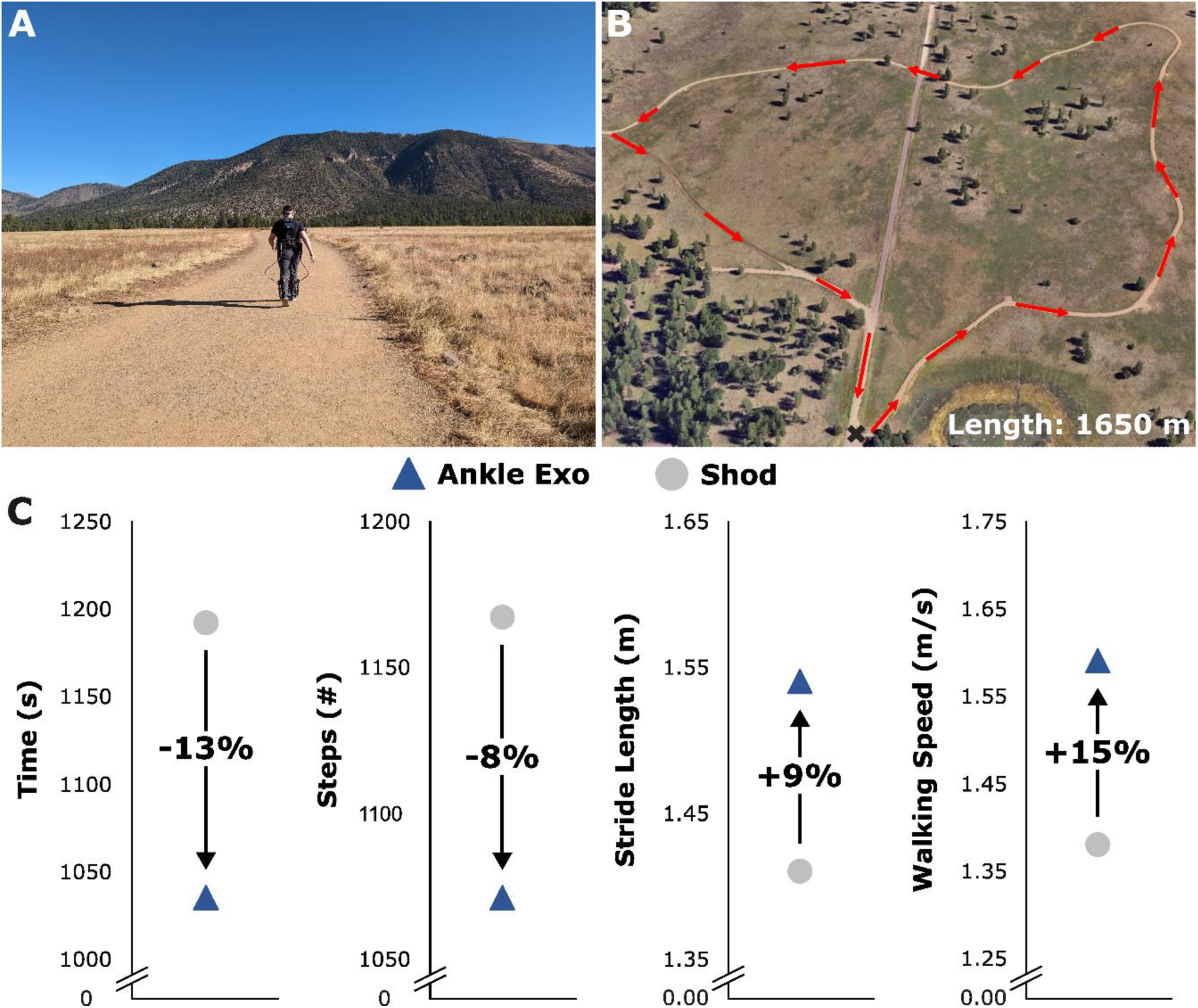
Ankle exoskeleton assistance increased mobility while walking in a real-world setting. **(A)** Participant walking around the outdoor course while receiving ankle exoskeleton assistance. **(B)** Outline of the outdoor course. **(C)** Experimental outcomes included: course completion time, number of steps taken with the right leg, average stride length, and average walking speed with (blue triangle) and without (grey circle) ankle exoskeleton assistance. The participant completed the loop in a shorter time by walking with a longer stride and faster walking speed while receiving ankle joint assistance.

In addition to completing outdoor testing, this same participant completed metabolic testing with and without the exoskeleton device during level indoor treadmill walking at a speed of 1.25 ms^-1^. Each walk lasted eight minutes from which the metabolic cost of transport during the last two minutes was utilized as a steady-state comparison between the two conditions. The participant saw an 8% reduction in the metabolic cost of transport while receiving ankle exoskeleton assistance, compared to walking without the exoskeleton (**Figure 5B**; Shod: 2.19 Jkg^-1^m^-1^; Exo: 2.02 Jkg^-1^m^-1^).

### Hip-&-Ankle – Load Carriage

To test the utility of the device while configured for simultaneous hip-and-ankle assistance, we had two individuals (**Supplemental Table 1**; P4 & P5) walk on a treadmill while wearing a weighted vest with a moderate sized load (22.5 %bodyweight). Each user walked for eight minutes in each configuration while metabolic data was collected via indirect calorimetry. The metabolic cost of transport was calculated for the last two minutes of each condition to enable comparison between the approaches. Both participants had reductions in the metabolic cost of transport with combined hip-and-ankle assistance (vs. no device) while walking on a treadmill carrying 22.5% of their bodyweight (**Figure 5C**). P4 had an 8% reduction in the metabolic cost of transport (Shod: 2.68 Jkg^-1^m^-1^; Exo: 2.46 Jkg^-1^m^-1^), compared to the shod condition, while P5 displayed an 18% reduction (Shod: 2.59 Jkg^-1^m^-1^; Exo: 2.13 Jkg^-1^m^-1^).

### Elbow – Weightlifting

With the device configured to provide elbow flexion assistance, two individuals (**Supplemental Table 1**; P6 & P7) performed weight curls with a 19.5 kg object until they reached fatigue and could not continue. A metronome set to sixty beats per minute was used to prompt users to raise and lower the weighted object (leading to a full cycle every two seconds) with a full repetition being considered one complete cycle from the down position back to the down position (**Figure 7A**). Prior to lifting, surface electromyography (EMG) electrodes were placed on the short head of the biceps brachii of each user’s dominant arm to record muscle activity during the task. Both participants saw large increases in the number of repetitions they could perform when using the elbow exoskeleton (**Figure 7B**). P6 increased their number of repetitions from 7 to 14 when going from no exoskeleton assistance to exoskeleton assistance (+100%), while P7 increased their number of repetitions from 6 to 22 (+267%). These increases in repetitions were supported by the EMG data as P6 had a 35% decrease in peak biceps activity and P7 had a 57% decrease in peak activity. Together, these indicate that elbow assistance during weightlifting may significantly increase endurance of the user which could suggest that utilizing this tool in a physically demanding workplace environment may increase productivity and/or decrease risk of injury. More extensive testing (n = 9) with this configuration has supported these findings with other users demonstrating reduced fatigue during lifting (the same task described here) and during more complex tasks like in a simulated workplace environment where they moved several 10 kg boxes from one shelf to another (*36*).

**Fig 7.**
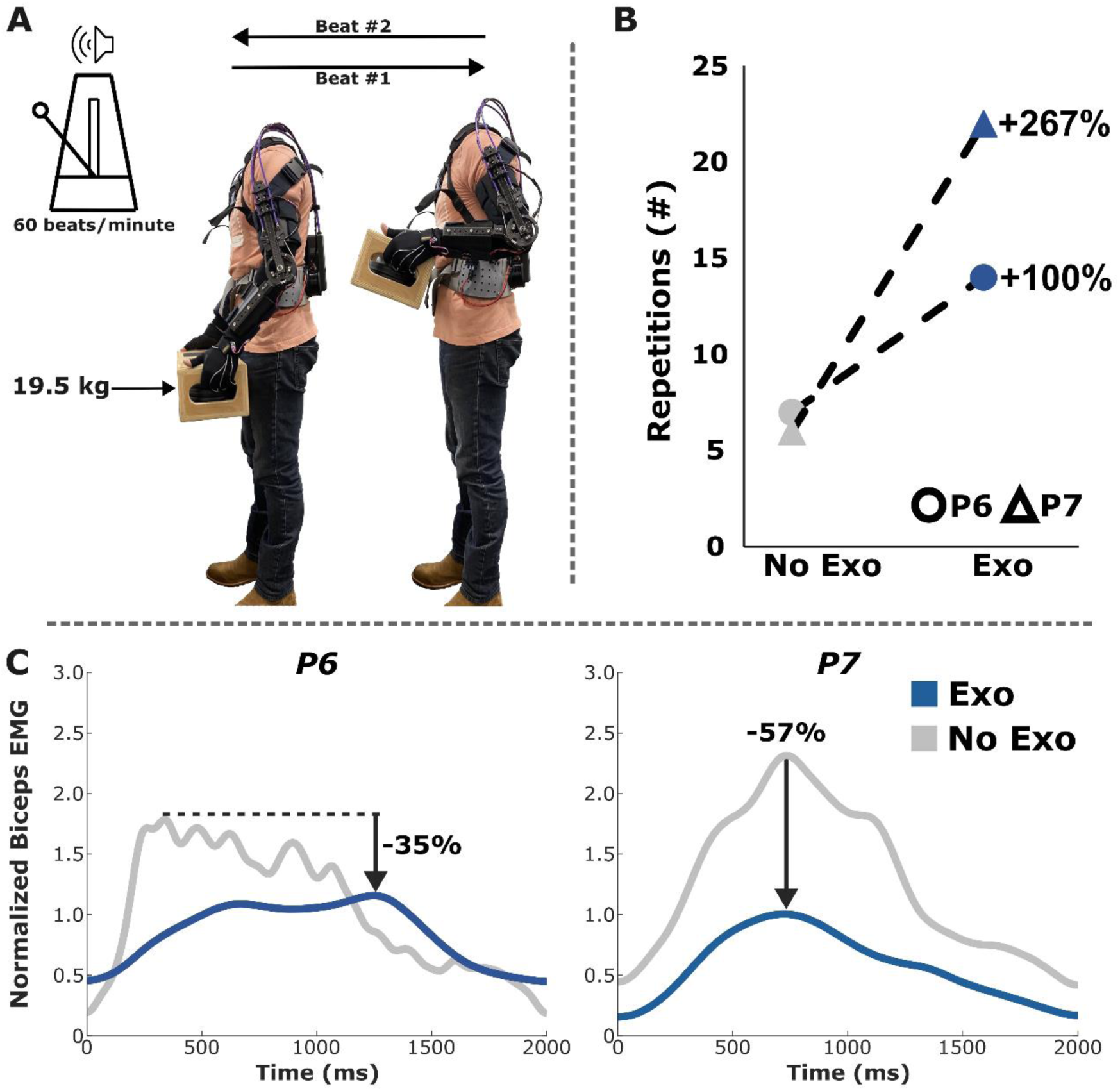
Elbow assistance during weighted lifting until fatigue resulted in decreased fatigue in users. **(A)** Experimental protocol, two users lifted a 19.5 kg box in time with a metronome while EMG activity was recorded from the short head of the biceps brachii of their dominant arm. **(B)** The number of repetitions increased by at least 100% for both users when receiving elbow assistance (blue) compared to not wearing the device (grey). **(C)** Similarly, large decreases in peak biceps EMG were detected with elbow joint assistance (vs. without assistance).

## DISCUSSION

This article describes the design and validation of a fully open-source exoskeleton system aimed at growing the field of wearable robotics. Importantly, this tool can be used as presented or as a starting point to be modified for the user’s own goals and interests. Here we outlined the design of the core-components to the system including the software framework, the electrical architecture, and its multiple hardware configurations and evaluated the device’s function under these configurations.

The most important feature of the system is its modularity. In particular, we believe that the presentation of a fully open-source and modular software system to control and operate exoskeletons is a unique contribution to the field as, to the best of our knowledge, most groups have provided little insight or access to this sub-component of their systems.

The creation of a software structure, particularly one that is adaptable to various user’s needs, is a time-consuming and tedious process that often requires significant troubleshooting and revision before productive research can be performed. For research groups who have iterated on years of prior closed-source work, this is not much of a barrier, but for those looking to get started in the field and those without extensive expertise, the design of this sub-component can be a daunting challenge that ultimately delays or prevents many from contributing to the field. Our hope is that those interested in pursuing lines of exoskeleton research can use this software, and the other tools that are a part of this system, to begin exploring new and unique applications of these devices. Simultaneously, we hope others who do have experience with exoskeletons can use this as an untethered testbed to begin to explore new hardware and control approaches that take these devices outside of the laboratory setting. Our long-term vision for OpenExo is for researchers from across the community to contribute to the growth and evolution of this system by sharing their hardware and control approaches developed with the system for all to access and test; ultimately accelerating the growth of the field, enhancing the reproducibility of findings, and moving us closer to a wearable future.

We tested the capabilities of the device across multiple different hardware configurations. When configured for open-loop, direct-drive hip assistance individuals saw metabolic improvements while walking on a 7.5^◦^ incline, suggesting that this device is capable of positively aiding movement during this task. These reductions in metabolic costs are consistent with prior work from our group and others, which suggests benefits could range from ∼10-15% while receiving hip assistance on an inclined surface (*22*, *41*). Similar metabolic benefits have been found with hip assistance during level ground walking (*37*, *42–44*) and running (*45*), as well as stair climbing (*46*) activities across both unimpaired and impaired populations. When walking indoors with ankle plantarflexion assistance we saw a metabolic improvement (vs. no exoskeleton) comparable to what others have found across a variety of tasks and populations (*29*, *37*, *47–49*). While the level of metabolic benefit across these configurations are largely similar, the design and control approaches to achieve these benefits vary wildly, limiting our understanding of what does and does not contribute to these improved responses. Future work by the community needs to refine our knowledge by isolating the design and control approaches that best improve performance under these configurations. When assisting both the hip and ankle simultaneously while users carried 22.2% of their bodyweight in additional load, we observed 8% and 18% reductions in the metabolic cost of walking. Fewer studies have examined multi-joint assistance, compared to single-joint devices, as multi-joint devices typically incur a large mass penalty due to the significant hardware needed to achieve this type of assistance. Those who have investigated multi-joint devices have overcome this challenge by utilizing tethered systems that offload a significant portion of the mass at the cost of being confined to the laboratory (*19*, *37*) or by exploring different hardware approaches such as soft exoskeletons (*50*, *51*) and/or combined passive and active systems (*52*). Preliminary work in the multi-joint domain suggests that they may be able to far exceed the capabilities of single-joint devices, with some finding that combined hip-knee-and-ankle assistance could reduce a user’s metabolic cost by upwards of 50% (*37*). The continued development of efficient hardware and controls approach for multi-joint devices is a key future direction for the field.

We had one user receive ankle joint assistance while on outdoor terrain and found that they had improved mobility, walking faster (+15%) with longer strides (+9%), compared to walking without the device. Most research with exoskeletons is performed in laboratory settings as it is easier to minimize potential confounding variables in these highly controlled environments. While these laboratory tests are valuable to understanding the potential of exoskeletons, more work must be done when utilizing these devices in their intended settings, such as on outdoor terrain, in clinical environments or even at home as the challenges associated with these environments (e.g., mixed-terrains and surfaces, differences in gait mechanics, and variable walking speeds) may dampen the utility of lab-based findings. For example, while the user in this study displayed improvements in walking speed when completing an outdoor, fixed-route, activity with ankle exoskeleton assistance, they also reported an increase in instability and a decrease in comfort as the application of power while walking on a dirt pathway seemed to result in altered traction between the user’s feet and the ground. This altered interaction between the foot and the walking surface would not have been detectable in a fixed laboratory environment or on paved outdoor surfaces and thus a key insight into how these devices interact with their users in this natural environment would have been missed. While researchers have begun exploring the potential for translating exoskeletons beyond the laboratory and into daily environments, they too have reported mixed results. Using an ankle exoskeleton with subject-specific control parameters informed by human-in-the-loop optimization, Slade et al. (*11*) found that users walked with a 9% increase in walking speed and were able to travel 17% further when compared to walking without the device in an outdoor environment. Tagoe et al. (*12*) found that ankle joint assistance could lead to metabolic improvements when assisting those with cerebral palsy on a mixed-terrain outdoor course. In contrast, MacLean and Ferris (*53*), who examined knee exoskeleton assistance during loaded incline walking in the lab and on mixed-terrain outdoor settings, only saw a benefit of assistance during indoor assistance. They hypothesized this could be a result of challenges associated with metabolic collections in outdoor environments and the intersubject variability associated with device usage including factors like subject-to-subject differences in anthropometrics, altered fit of the device, and the variability in how individuals’ fatigue and acclimate to devices. It is quite clear that outdoor environments present significant challenges to evaluating the viability of devices and control strategies when compared to the lab. One reason for this challenge could be the field’s overreliance on metabolic cost as a means for verifying the utility of devices. This measurement is often highly variable and thus is extremely dependent on using steady-state tasks to have a degree of validity. Unfortunately, mixed-terrain, real-world settings are often not conducive to steady-state tasks and thus make user’s energetics a relatively inaccurate and poor measure in these domains. Future work aimed at identifying new ways to evaluate device efficacy is paramount for the field to expand beyond the lab and into the real-world.

While lower-extremity devices have had a hard time moving beyond the lab, upper extremity devices have had more success given their task-oriented nature (*54*, *55*). In this study, we outlined a novel elbow exoskeleton operating on this open-source infrastructure. Using this configuration, we found that users experience benefits from elbow assistance, as they were able to increase the number of weightlifting repetitions (+100% and +267%) while seeing corresponding reductions in biceps muscle activity (−35% and −57%). A full breakdown of this device, its novel control paradigm, and its utility during more realistic labor tasks can be found in Colley et al. (*36*). Previous work exploring other upper extremity targets like the shoulder and back have demonstrated benefits that could be applied successfully in ergonomic settings (*14*, *17*, *20*, *21*). The elbow joint has been targeted less often (*55*), with only a handful of previous studies having explored untethered devices, with most being passive in nature (*56*). These previous attempts at assisting this joint have lacked autonomous usability and have generally been tested during only static tasks (*57–60*). The elbow configuration developed on this open-source system is among the first to provide fully autonomous usage while also enabling dynamic work to be performed. Little work has been done exploring the utility of combining upper and lower extremity systems, despite their obvious ergonomic potential. Future work exploring this domain and its uses for real-world applications (e.g., military, physically demanding workplace environments) may be the shortest route to integrating exoskeletons into our everyday lives.

There are several promising future directions for OpenExo. First, we will continue to update our documentation for the system as we develop new components for it and receive feedback from the community. Second, the system is currently limited by its battery life (a common challenge with untethered, autonomous systems), especially when utilized in high-torque multi-joint applications. We have provided recommendations for other sources of power to the system that may increase operational lifespan and will continue to explore and update users on potential solutions to increase operational duration as new research developments are made. Third, we will continue to update and optimize the companion python API and GUI available to users to help control the device. To date, it has several features, such as the ability to record and store data, plotting, and updating controllers and parameters in real-time but lacks other core components like exo-informed biofeedback (*61*). Further expanding this application to include some of these core features as well as other state-of-the-art research capabilities (e.g., deep learning (*62*) and human-in-the-loop approaches (*49*)) is a high priority. Finally, the experimental validation of the configurations presented in this paper contained small sample sizes. While larger sample sizes would increase confidence in the findings, our primary goal was to highlight the modularity of the system and provide examples of its potential benefits during typical research tasks. Given the open-source nature of the device, we believe it represents a promising solution to rectify the small-sample size nature of most exoskeleton studies in literature. Specifically, this standardized system could enable increased collaboration and large multi-site studies exploring applications of exoskeletons, something that has previously evaded the community.

In summary, we believe OpenExo will facilitate future directions for the field. Several excellent reviews have summarized the current state of research related to wearable robotic devices (*5*, *63*, *64*). This includes highlighting work centered on tethered systems and their use as laboratory-based testbeds to explore novel control schemes (*19*, *37*), the combination of passive and active designs to enhance the ability and lifespan of devices (*52*, *65*, *66*), and on the end-user themselves; trying to understand how individuals acclimate (*67*) and respond to devices (*68*), and what differentiates who does and does not respond to certain devices. There has also been a large push by the field to incorporate aspects of machine learning and artificial intelligence to expand the capabilities of these devices, especially on the control end of these applications. Critical future developments include the continued expansion of exoskeleton usage beyond the laboratory setting, including hybrid devices to assist or rehabilitate individuals with impairments or injuries in the clinic and at home, and the continued exploration of new and exciting ways to understand how we can best evaluate these devices, their impact on us, and how they can best serve us in society. Our hope is that this OpenExo system will present opportunities for those with valuable expertise outside the domain of exoskeletons to lend their experiences to help advance some of these research directions. For example, those who have a background in clothing design/fashion, who would never have the extensive expertise needed to build up an exoskeleton, may be able to use this system to test different approaches to maximize the comfort and fit of these devices without the worry of designing software or electrical components. Similarly, computer scientists who may lack the technical expertise needed to design and construct systems but who do have expert knowledge in emerging domains like artificial intelligence and machine learning may be able to incorporate their knowledge and make contributions to developing highly adaptable exoskeletons capable of usage in any setting. This could also include experts from other domains such as material science, renewable energy and smart batteries, thermo-fluid systems, novel sensing and actuator development, and medicine. Like all research, it will be a team effort to advance these devices for use in our daily lives. Finally, we hope that this system will serve as a valuable educational tool for any who are interested in starting work in the field of wearable robotics, as the continued inspiration and education of new researchers with diverse life experiences and prospectives is an important building block to the development of fully realized devices to change the lives of those in need.

## MATERIALS AND METHODS

OpenExo consists of four subsystems: (1) electronics, (2) software, (3) hardware, and (4) control. Each of these subsystems will be detailed independently, but for the sake of brevity we will only highlight the design and control approaches in this section. Information on the electronics and software subsystems as well as the methodology for the benchtop and experimental validation can be found in the **Supplementary Material**. More detailed information can be found on the system’s dedicated website (theopenexo.org).

### Exoskeleton Design

The system was designed to maximize flexibility and allow for multiple configurations. Each configuration is centered around the same base human-robot interface in the form of a waist belt containing the electrical components and each operates on the previously described software structure. While this waist belt assembly is consistent between devices, motor mounting strategies differ based on the desired task (i.e., hip vs. ankle vs. elbow).

### Waist Belt

The waist belt is the core connection between the user and the robotic hardware. Thus, finding a sturdy but comfortable belt that can accommodate multiple motors at a time (up to four) was a high priority. We settled on an adjustable hip belt used for outdoor recreational activities (Osprey IsoForm 4), which has adequate padding for comfort. Four sizes (extra small, small, medium, large) were utilized to provide flexibility to conform to the wearer. Each belt was modified by attaching a custom designed 3D printed mount for the waist assembly containing the PCB and battery. A 22.2v, 1800 mAh Li-Ion battery (HRB) was utilized throughout the study (**Supplemental Figure 2**).

### Hip Device

#### Hardware

We designed a bilateral, direct-drive untethered hip configuration capable of providing flexion and extension assistance during a variety of tasks such as level and incline walking (**Figure 2A**). The main design goals were to create a device that (1) is simple to construct, (2) could provide relevant magnitudes of assistance while staying lightweight, (3) is low-cost, and (4) can adjust to a broad range of anatomical geometries. To accomplish this, the waist belt assembly described above was modified to allow for the placement of motors in-line with the hip-joint center. Our design consisted of three main components: (1) the modified waist belt, (2) the belt-motor connection, and (3) the motor-thigh assembly. The waist belt was modified by adding mounting plates on each side of the belt in line with the hips (**Figure 2A**). A grid of equally spaced M5 clearance holes were drilled into the mounting plates to allow for flexibility in placement of the belt-motor connection to account for anatomical variability from person-to-person. The belt-motor connection consisted of an aluminum abduction/adduction hinge capable of facilitating out of plane motion during walking and carbon fiber brackets to interface the motor with the hinge. We used AK60-6v1.1 motors (CubeMars; Nanchang, Jiangxi, China) with each motor capable of both hip flexion and extension assistance. These versions of the AK series motors were selected due to the relatively lower torques required of hip assistance (compared to the ankle joint) and their lower mass compared to more powerful alternatives (e.g., AK80-9). These motors were connected to a carbon fiber upright via a custom-designed interface consisting of carbon fiber and 3D printed onyx materials (carbon-reinforced nylon; Markforged; **Figure 2A: Motor Mount Assembly**). This interface directly connects to the carbon fiber upright, causing the upright to actuate in the direction that the motor spun. Thigh cuffs were designed to slide onto the carbon fiber upright to help facilitate torque transmission from the motor-upright configuration to the user. These cuffs were designed to conform to the shape of an individual’s leg, were outfitted with padding to provide comfort, and included an adjustable strap mechanism to ensure good fit with the user regardless of thigh size. Different sized thigh cuffs were designed with the goal of being easily swappable depending on the size of the user.

Interfacing with the hardware located at the hips and thighs are bilateral heel and toe FSRs placed underneath the insoles of the wearer’s shoes. These measure the relative force placed on each aspect of the foot and allow for control paradigms based on estimates of heel and toe contact with the ground, such as gait-phased based control schemes. The total mass of this configuration is 2.9 kg.

#### Control

Our primary mode of control for this hip device was a variation of a control scheme developed by Bryan et al. (*19*) and Franks et al. (*37*) (**Figure 8A**). Briefly, heel and toe FSRs were used to estimate the gait cycle of the user. This was done by measuring the time duration from successive heel-strikes. Then, the gait phase was estimated by calculating the current time from the last heel-strike relative to the expected duration based on the average of the previous three steps. Users could then prescribe hip torques based on the percentage of the gait cycle. As outlined in Bryan et al. (*19*) and Franks et al. (*37*), the starting point of hip assistance was shifted to 84% of the gait cycle to prevent discontinuities related to the need for hip extension assistance during heel strike. Based on this, the user could define several controller parameters such as the magnitude and percent gait cycle of peak flexion and extension torques as well as the rise/fall time, in terms of percent gait cycle, to reach those peaks. These points were connected using splines to facilitate a smooth transition during assistance. For timing-based measures (percent gait where peaks occurred and the rise/fall times to get there), we elected to use the average parameter values reported by Franks et al. (*37*) as they found that optimized torque timing was relatively consistent between subjects. Thus, for the purposes of this study, only flexion and extension torque magnitudes were varied on a subject-by-subject basis. This controller, and others, are available for use as part of our software architecture.

**Fig. 8.**
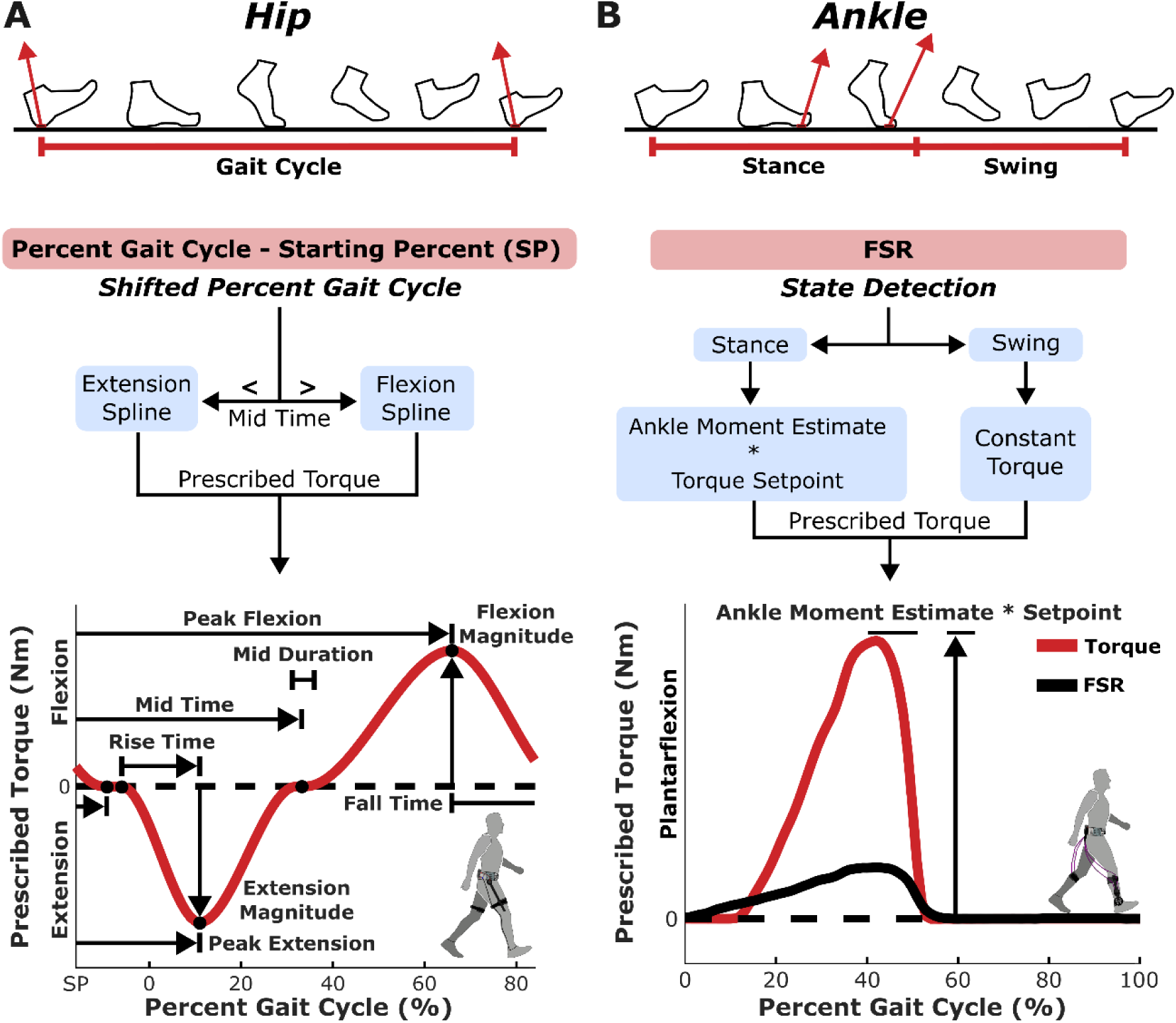
Control profiles utilized for the hip and ankle configurations. **(A)** The hip exoskeleton is controlled based on an estimate of the user’s current position in their gait cycle (Percent Gait Cycle) as calculated by the time since their last heel strike relative to their average time from heel strike to heel strike. The prescribed torque profile is then applied via extension and flexion spline curves that are based on a user defined shift in percent gait (Shifted Percent Gait Cycle, to avoid discontinuities at heel-strike) and user-defined parameters (i.e., rise time, extension magnitude, peak flexion, etc.). This controller is based on the works of Bryan et al. (*19*) and Franks et al. (*37*). **(B)** The ankle exoskeleton is controlled via a previously described proportional joint moment controller (*29*, *38*). This operates by detecting the current state of the user (stance or swing) based on a FSR placed at the forefoot. If the user is in stance, an estimate of the ankle moment is determined based on a previously validated regression equation that takes a normalized FSR value as its main input. This moment is then scaled based on a user defined setpoint resulting in an assistive torque profile that follows the shape of the user’s estimated ankle moment but with a torque that is only a percentage of that moment. If in swing, a constant user defined torque can be supplied if desired.

### Ankle Device

#### Hardware

Our untethered, bi-directional sagittal-plane ankle configuration was similar in design to our previously reported ankle device (*29*) (**Figure 2B**). Briefly, motors mounted at the waist actuated a pulley located at the user’s ankle via a cable-chain interface which, in-turn, created sagittal plane movement (either plantarflexion or dorsiflexion) for a carbon fiber footplate that could slip into any user’s shoes. The ankle assembly portion of the device was identical to the previously reported device. To summarize, a single-degree of freedom (sagittal-plane rotation) pulley interacted with a user via carbon-fiber footplates and custom-designed calf cuffs. To facilitate a variety of user sizes, multiple sets of footplate and calf cuff sizes were created with the ability to quickly be swapped out with the rest of the device assembly. This pulley was actuated by steel cables driven by the motors located at the waist. In-line with the pulley were custom-designed torque and angle sensors which were utilized for low- and high-level control schemes, respectively. For more information on the torque and angle sensors and their validation, we point you to (*29*). The pulley, which was placed in a carbon-fiber upright that interfaced with the user via the custom-designed calf cuffs, formed a 4.5 gear reduction ratio relative to the motor. The device was designed to minimize distal mass and lateral protrusion in order to reduce the metabolic burden and potential for out-of-plane bending moments/environmental collisions, respectively. To facilitate high-level control schemes, FSRs were placed on each footplate near the ball of the foot (under the distal head of the first metatarsal).

While the ankle-assembly remained nearly identical to our previous design (*29*), the waist-mounted Bowden-cable transmission portion of the device was redesigned. This was done for two reasons: (1) to accommodate new motors and (2) to facilitate the modular nature of the designed system. Motors were placed externally to the electrical casing to allow for rapid switching to a new device configuration (e.g., hip actuation only). 3D printed interfaces were designed to attach the motors to the waist belt (**Figure 2B: Motor Mount Assembly**). To further enhance modularity, these 3D printed interfaces were designed to facilitate multiple versions of the AK series motors (e.g., AK80-9, AK60-6v1.1, AK70-10). For the sake of brevity and clarity, all references to the ankle device will be in its AK80-9 configuration. A custom-designed 26 mm sprocket was fastened onto each motor to turn chains within a 3D printed cartridge. This, in turn, led to the actuation of the cable transmission system that rotated the ankle pulley, thus translating torque from the motor to the wearer. The steel cables interfaced with the chain via custom-designed aluminum interfaces. These steel cables were passed through Bowden sheaths to guide them down to the ankle assembly. At the bottom of the 3D printed cartridge, we designed and implemented a strain-relief system to minimize high-strains where the Bowden sheathes exited the cartridge. This system consisted of a 3D printed onyx-casing that interfaced with 3D printed thermoplastic polyurethane (TPU) inserts that housed the Bowden sheaths (**Figure 2B**). The design of the waist-belt Bowden-cable transmission system is available for others to replicate as part of this open-source system (**Supplemental Figure 1**).

The total mass of the ankle configuration, when utilizing AK80-9 motors and when sized to a 6-foot-tall adult male, is 3.9 kg. This likely represents the largest weight for the system, as the motors (and the associated cartridges), cable-transmission system (length of steel cables and Bowden tubes), and the custom-designed footplates and calf-cuffs were all at their largest.

#### Control

The ankle configuration was controlled using a previously described proportional joint moment controller (PJMC) developed by our research group (*29*, *38*) (**Figure 8B**). Briefly, we used FSRs placed at the forefoot to detect the stance and swing phase transitions of gait. During stance, the controller generates an adaptive plantarflexor torque based on a real-time estimate of the biological sagittal plane ankle moment. Users specify a setpoint to limit the maximum torque output and the adaptive plantarflexor torque is scaled based on this setpoint. This results in an assistive torque profile that follows the shape of the user’s ankle joint moment but with a torque magnitude that is typically only a percentage of that moment. While in swing, the user has the option to prescribe a constant dorsiflexion torque to aid in toe clearance. Throughout, a low-level PD controller is used to facilitate closed-loop control to ensure the prescribed torques matched those measured from a torque transducer placed in-line with the sagittal plane of the ankle joint. The gains of the p and d terms were determined via manual tuning while a user walked on a treadmill, receiving exoskeleton assistance (P:28, D: 200).

### Hip-&-Ankle Device

#### Hardware & Control

The hip and ankle devices described above were combined into one system to provide multi-joint actuation (**Figure 2C**). The footplates of the ankle device were modified to include a heel FSR in addition to the toe FSRs already embedded within the plate. The same control schemes used in the single-joint configurations were utilized to provide multi-joint assistance.

### Elbow Device

#### Overview

To demonstrate the capacity and flexibility of the designed open-source framework, we also developed and constructed a bilateral, sagittal plane, elbow exoskeleton (*36*) (**Figure 2D**). A full description of the design of the hardware and control is outlined in Colley et al (*36*), here we will provide a general overview of this configuration. Briefly, using a similar Bowden-cable driven transmission as the one used for our ankle exoskeleton device, motors (AK60-6v1.1s) mounted at the waist actuated steel cables that rotated a pulley placed in-line with the elbow joint. This system interfaced with the user via custom-designed carbon fiber uprights and 3D printed forearm and upper-arm cuffs as well as an upper-torso harness. The cuffs, the harness, and the waist belt were all adjustable to fit a wide variety of user dimensions. To facilitate low-level control schemes, we placed a torque transducer in-line with the pulley and elbow to estimate sagittal plane elbow moments. To facilitate high-level control, small FSRs were placed on each hand at the proximal phalanges of the middle, ring, and little fingers (wired in parallel to create a single FSR signal) and on the palm at the base of the thumb. The device operated at zero torque (only applying torque to offset the resistance of the cable-driven system) when the FSR value was below a user-defined threshold and provided flexion assistance when it exceeded this threshold. A low-level PD controller was used to perform closed-loop control to ensure the prescribed torques matched those measured from the torque transducer.

## Supplementary Materials

**Supplemental Figure 1.**
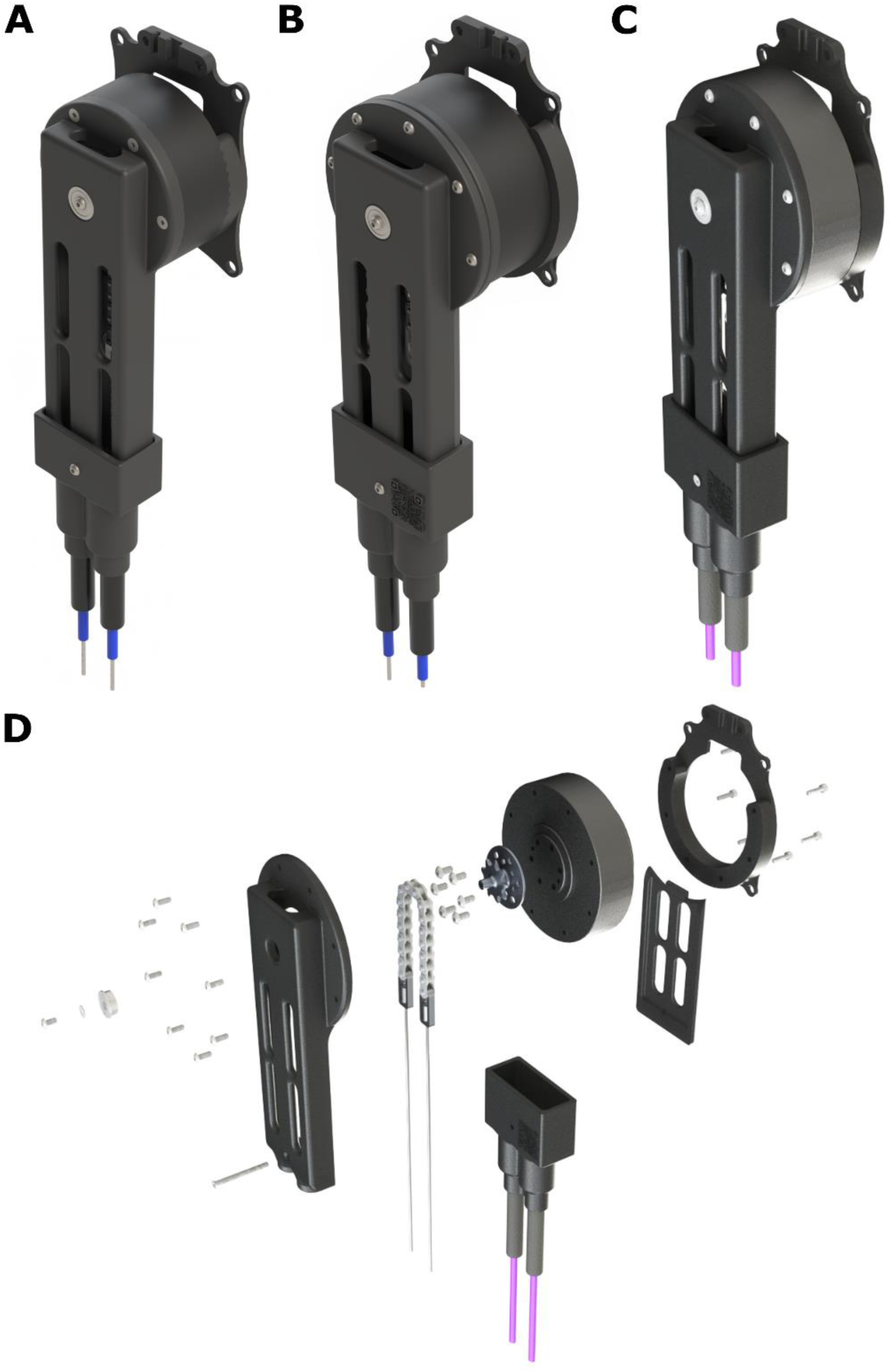
Open-Source Transmission Design. **(A)** AK60-6v1.1 motor, **(B)** AK70–10 motor, **(C)** AK80-9 motor, **(D)** Exploded view of AK80-9 motor assembly. A full breakdown of the required parts and an assembly guide are available within OpenExo’s documentation. The same transmission style has been designed to accommodate a variety of CubeMars AK-motors including the AK60v1.1, AK70, and AK80. The parts were designed to fit to the belt in the same manner, minimizing the amount of revision necessary to attach to the waist belt. The same belt-side transmission was used for both the ankle and elbow joints, with only the end-effector hardware differing between the two configurations.

**Supplemental Figure 2.**
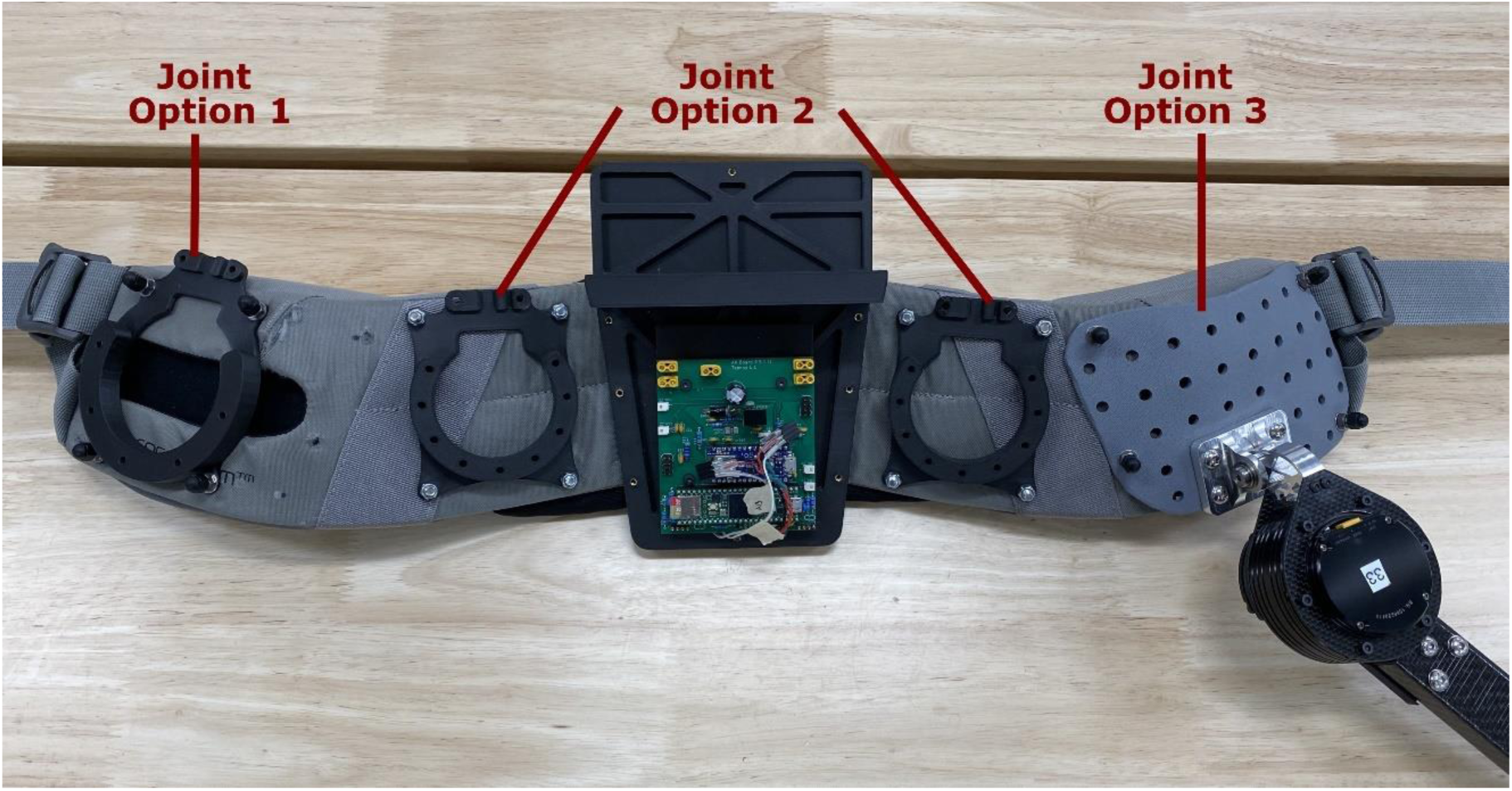
Modularity of the waist belt design. The waist belt contains an electronics and battery case containing the PCB (that can support the bilateral operation of two joints at a time) and the battery. The belt, however, is modular enough to be utilized in several configurations. The image above highlights how different orientations can be achieved on the belt. This can include two Bowden-transmission joints (Joint Option 1 & 2 in the image, with the right-side Joint Option 1 currently being covered by Joint Option 3) or one Bowden-transmission joint and one direct-drive joint (Joint Option 2 & 3 in the image, with the left side capable of taking a similar plate as the right to accommodate bilateral operation). Plausible configurations with the established joints could include single joint assistance (hip, ankle, elbow) or multi-joint assistance (hip-ankle, hip-elbow, ankle-elbow).

**Supplemental Figure 3.**
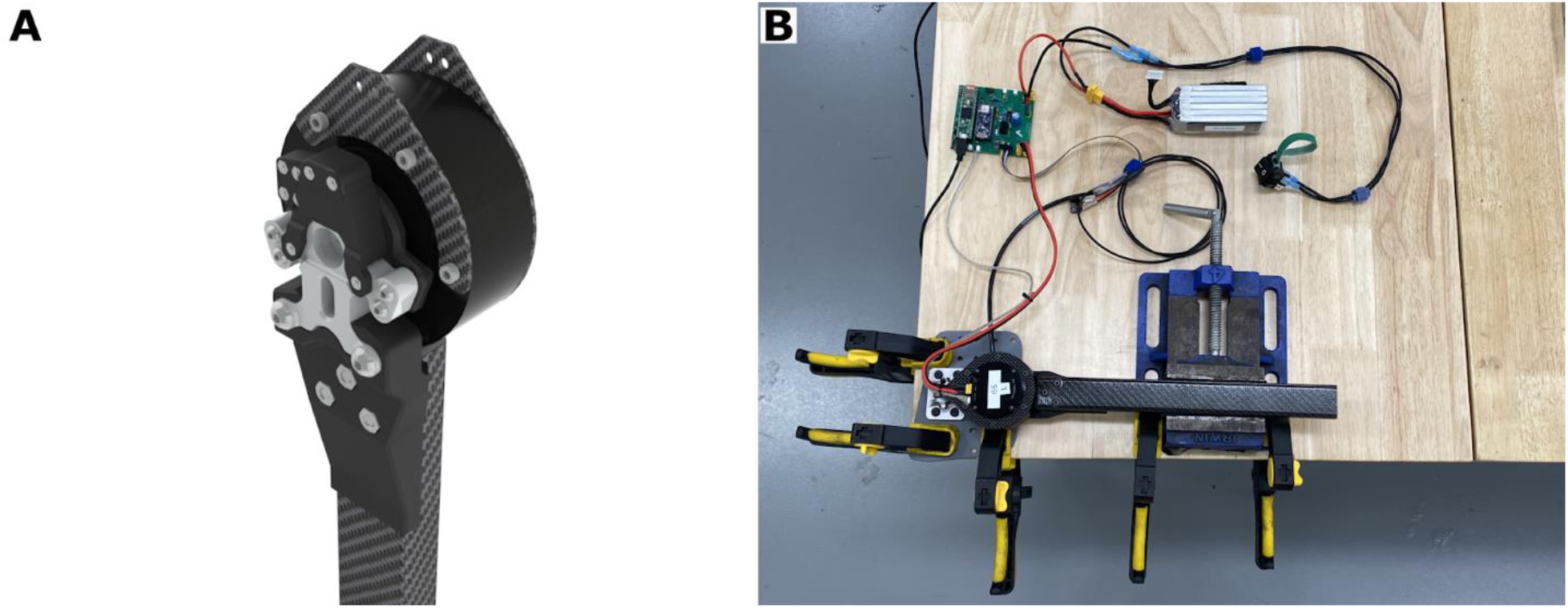
Experimental setup utilized to perform the engineering validation of the hip configuration. **(A)** Modified hip exoskeleton design incorporating a torque transducer. This modified design featured altered 3D printed components to link the motor to the carbon fiber upright while incorporating the torque sensor into these components. This version of the design was only used for the engineering validation work. **(B)** Image of benchtop testing setup. The motor-upright assembly was secured to a table via clamps and vices to prevent movement. The codebase was modified to include a new exoskeleton controller designed to apply step commands with user defined amplitudes, duration, and spacing (included as part of OpenExo’s software). A series of open-loop step commands were sent to the motor while recording the measured torque.

**Supplemental Table 1:** Participant Characteristics.

| Configuration | Condition | Participant | Sex | Age (years) | Mass (kg) | Height (cm) | Torque (Nm) | Walking Speed (m/s) |
| --- | --- | --- | --- | --- | --- | --- | --- | --- |
| **Hip** | Incline walking (7.5°) | P1 | M | 23 | 65.6 | 169.0 | Ext: 4.1<br>Flex: 3.4 | 1.0 |
|  |  | P2 | F | 63 | 50.6 | 167.5 | Ext: 4.1<br>Flex: 3.4 | 0.85 |
| **Ankle** | Outdoor | P3 | M | 28 | 93.0 | 182.0 | PF: 28 | Not Fixed |
|  | Indoor |  |  |  |  |  | PF: 28 | 1.25 |
| **Hip-&-Ankle** | Load carriage (22.5% BW) | P4 | M | 32 | 70.8 | 172 | Ext: 4.1<br>Flex: 3.4<br>PF: 28 | 1.25 |
|  |  | P5 | M | 37 | 71.1 | 173.5 | Ext: 5.5<br>Flex: 4.9<br>PF: 28 | 1.25 |
| **Elbow** | Weight lifting (19.5 kg) | P6 | F | 22 | 65.7 | 156.5 | Flex: 12 | NA |
|  |  | P7 | M | 23 | 79.1 | 171.8 | Flex: 12 | NA |

Note: Ext = Extension; Flex = Flexion; PF = Plantarflexion

## Supplemental Information - Figure 2 A&B Labels

### Hip Configuration

The exploded view of the hip assembly contains the following components: (1) carbon fiber upright, (2) 3D printed screw mount, (3) 3D printed shell for the motor-to-upright connector, (4) carbon fiber motor-to-upright connector, (5) carbon fiber motor space, (6) inner carbon fiber motor bracket, (7) AK60-6v1.1 motor, (8) aluminum abduction/adduction block, (9) outer carbon fiber motor bracket.

### Ankle Configuration

The exploded view of the ankle assembly contains the following components: (1) Bowden sheath, (2) TPU strain relief, (3) strain relief casing, (4) outer motor cartridge, (5) steel cables, (6) aluminum cable-chain interface, (7) chain, (8) aluminum sprocket, (9) AK80-9 motor, (10) back motor cartridge, (11) Motor-to-belt connector.

## Supplemental Information – Additional Methods

### Electrical Architecture: Microcontrollers, Actuators, Sensors, and Printed Circuit Board

#### Microcontrollers

We developed a simplified electrical architecture, which featured unused or networked electrical connections to support multiple potential joint configurations. At the center of this architecture are two microcontrollers: (1) a Teensy 4.1 Development Board which was used to perform tasks such as data acquisition, control, and actuator communication, and (2) an Arduino Nano BLE 33 which was responsible for communication with a companion application via Bluetooth Low Energy (BLE). Two microcontrollers were selected, rather than one, to increase device speed. The main goal was to find one microcontroller for exoskeleton control and one for Bluetooth communication. The Teensy was selected for the exoskeleton control due to its fast clock and good analog-to-digital converter, while the Nano was chosen for its Bluetooth capabilities. Both microcontrollers can directly interface with Arduino to facilitate easy usage of the developed software system.

#### Actuators

We designed this system around CubeMars’ AK-series motors (Nanchang, China) which are powerful enough to provide torques for larger adult participants and during more challenging tasks such as multi-terrain walking (e.g., stairs, inclines, declines…). Our electrical architecture, software, and hardware were all designed to be able to interface with different versions of these motors to provide the researcher with maximum flexibility. All real-time communication with these motors is performed using the CAN protocol, which is a bus-based communication protocol that relies on differential signaling for noise rejection. It should be noted that our software system is robust enough to integrate different motors other than the CAN-based ones utilized currently. We have developed simple guidelines for users to follow if interested in adding a new motor to the system.

#### Sensors

The system was designed to allow for multiple, independent, sensors. Commonly used sensors for our devices include a proprietary, strain-based, torque transducer (*29*) to estimate applied torque at a joint to facilitate low-level closed-loop control, force sensitive resistors (FSRs) placed around the ball of the foot and heel to enable high-level control, and hall effect magnetic angle encoders to estimate joint position and velocity. To account for variance in how the device fits users, which will impact the signals and estimates from each of these sensors, we developed range-based calibration protocols to be employed after donning the device to ensure consistency. Each sensor is available to use but not a required component for the designed architecture to function properly (i.e., if no angle sensor is present, that wouldn’t prevent the device from functioning if the high-level control did not require it). A step-by-step guide is available as part of our software structure should researchers have interest in adding additional sensors to their device.

#### Printed Circuit Board

We designed a printed circuit board (PCB) that was used to power and communicate with the actuators and to condition analog signals from external sensors (e.g., torque transducers, FSRs, angle sensors, etc.) while directly interfacing with the microcontrollers (**Figure 1C**). This included ports to power and communicate with up to four actuators at once (allowing for bilateral multi-joint applications), as well as pins to interface with multiple sensors (e.g., simultaneous usage of bilateral torque sensors and FSRs). Documentation breaking down the components of the PCB and guides to modifying the software to incorporate new PCBs are included to help facilitate modification of the system to meet a user’s needs.

### Software: A Modular Code Base for System Flexibility

#### Overview

The chief source of modularity in our open-source system comes from the software architecture. Our software was developed using a combination of C++ and Arduino languages, with the principle of inheritance-based polymorphism (realized through parent-child classes and abstract classes) used to achieve a high-degree of modularity and reduce redundancy. **Figure 1B** outlines the structure of the software. We designed the software using a hierarchical structure in order to have a logical and easy to follow form. We also used a parallel structure to facilitate the dual usage of microcontrollers. Thus, there were essentially two components to the code: (1) the main classes for handling computation (e.g., Exo, Side, etc.) and (2) classes designed to store the data and parameters used in control of the system (e.g., ExoData, SideData). The structure of these two components mirrored each other with the data classes (e.g., ExoData) inheriting their interfaces from their computational (e.g., Exo) counterparts via abstract classes. This was done so that the classes responsible for the data (e.g., SideData) could be mirrored onto the Arduino Nano microcontroller in order to interface with the Bluetooth communication without the need to mirror the components of the software used for the primary exoskeleton control (thus improving operating speed).

#### Structure

The exoskeleton device was designed to interface with a companion python application (**Figure 1B: Python API**). This device communicates with the system through the Arduino Nano microcontroller via BLE. This microcontroller handles all the Bluetooth communication of the device and contains a mirrored copy of the exoskeleton data which originates from the Teensy microcontroller, with data communication between the two microcontrollers occurring via universal asynchronous receiver/transmitter (UART). The Teensy microcontroller handles most of the computation that occurs within the system. This includes configuring the device via user defined parameters stored in a SD card that slots into the Teensy (**Figure 1B: SD Card**), reading the primary sensors associated with the system (e.g., FSRs, torque transducers; **Figure 1B: Sensors**), computing the primary outputs of the system (i.e., motor commands based on the joint configuration and desired controller utilized), and communicating the primary outputs with the relevant hardware (i.e., the motors; **Figure 1B: Motors**).

#### SD Card

An SD card is used to store information regarding the device configuration as well as to store default values for the parameters associated with the controllers for each joint configuration. This SD card is slotted into the Teensy and the software operating on the microcontroller then accesses the stored data to get information about the desired configuration of the device. This includes PCB information (i.e., version information which changes pin readouts associated with both microcontrollers), which joints are used (i.e., hips, ankles, hips and ankles, knee, elbow), the sides used (i.e., unilateral or bilateral), transmission gearing ratios associated with each joint, default controller information (i.e., which controller should be default for the given configuration), sensor usage information, and directionality information (i.e., whether the calculated motor commands should be flipped to match a specific direction). Additionally, parameter information for each controller for each joint is stored on the SD card; this may include information such as max torque setpoint for a specific controller, information about when to begin torque onset, and PID gains for closed-loop control. These parameters are the default settings for the specified controller and can be accessed and changed in real-time via the companion application. Step-by-step guides are available to assist users in adding new joint configurations or controllers/controller parameters to the software to help facilitate researcher usage.

#### A Python GUI for Exoskeleton Control

A companion python application was developed to facilitate easy usage of the exoskeleton device. Core functions of this software include connecting to the device, calibrating the sensors, starting an active trial, updating controller parameters in real-time, and plotting. Upon the completion of the trial, data from up to ten variables of interest can be saved and exported as a .csv file to enable researchers to analyze the data and verify the accuracy of their intended control paradigm. Like the base software for the exoskeleton, this companion application is open source and available for researchers to download and modify to match their needs.

### Engineering Validation

#### Hip Configuration

##### Benchtop Testing

To characterize the responsiveness of the direct-drive hip configuration, a benchtop step-response test was performed. To confirm the accuracy of the device’s response, the design of the hip was modified to include an in-line torque transducer between the waist-mounted motor and the carbon fiber upright (**Supplemental Figure 3A**). This torque transducer was only used to verify the torque being produced at the motor and was not used for any low-level control approaches (e.g., PID control); that is, all tests performed with this modified design were under open-loop control. The mounting plate containing the hip assembly was removed from the waist belts and secured to a table via external clamps. The carbon-fiber upright of the assembly was also secured to the table via a vice to prevent movement (**Supplemental Figure 3B**). A step response controller was developed in the previously described software and allowed users to specify the amplitude, duration, and spacing of the step commands applied by the system. For benchtop testing, the peak amplitude of the step command was set to 6 Nm (0.5 Nm higher than the maximum assisted torque from the experimental validation). A total of five commands were sent, each lasting two seconds with two seconds of spacing between each command. The duration, prescribed magnitude, and the measured torque were recorded for each step command and averaged across all five steps. From this data, the rise time, defined as the time it took for the measured torque to go from 10% of the prescribed torque to 90% of the prescribed torque, and the percent overshoot/undershoot, defined as the difference between the average measured torque at the peak of the step and the prescribed torque, were determined. Residual motor vibration while applying high torques to a rigidly fixed system caused a high degree of low-magnitude oscillation when at the maximum torque setpoint for this test, resulting in unrealistic overshoots compared to how the device would respond when worn by a user. To reduce this noise and more accurately capture the performance of the system when worn by a user, a real-time exponentially weighted moving average was applied to the measured torque once it reached its maximum torque setpoint to help smooth out this noise. Importantly, this had no impact on the calculation of the rise time (due to the lack of applied filter before reaching the setpoint) and minimal impact on the average overshoot of the torque setpoint (as the noise was relatively uniform around this mean).

##### Torque Tracking

To characterize the accuracy of the previously described hip controller, the modified hip design featuring the in-line torque transducer was used during walking to record the measured torque experienced by the user. Despite the presence of the torque transducer, the user walked with open loop control to provide the most accurate representation of how the hip device would function (without the in-line transducer). That is, the transducer was only present to verify the accuracy of the prescribed open loop controller rather than to help influence the control. The individual walked at the most commonly reported maximum flexion and extension torque magnitudes from our experimental validation (Flexion: 3.4 Nm; Extension: 4.1 Nm). While they walked, the estimated percent gait cycle, the prescribed torque, and the measured torque were recorded. The user walked in the device for two minutes and the averages and standard deviations of the three measures of interest were determined over the course of the trial. The accuracy of the prescribed controller was characterized by the average root mean squared error (RMSE) between the prescribed and measured torques.

#### Duration Testing

A duration test was performed with a user walking at their maximum comfortable hip assistance (Flexion: 5 Nm, Extension: 5.5 Nm, Treadmill speed: 1.25 ms^-1^; as determined from experimental validation). As they walked the motor temperature for both the left and right motors and the total battery voltage (from the 22.2v, 1800 mAh Li-Ion battery) was recorded every minute via an infrared thermometer (Etekcity Lasergrip 1080) and LiPo Battery Voltage Tester, respectively. The test was concluded once the motors shut off due to high temperature or once the battery voltage reached the manufacturer’s minimum voltage recommendation (3.7 V/cell = 22.0 V total).

### Ankle Configuration

#### Benchtop Testing

Like the hip configuration, we tested the responsiveness of the ankle configuration via benchtop step-response testing. This consisted of having a user don one of the legs of the exoskeleton device and having the footplate and carbon fiber upright firmly fixed to their lower leg while the user was seated with their leg bent 90° at the knee. The same step response controller described previously was then applied with the maximum torque amplitude set to 28 Nm (the experimentally determined maximum comfortable plantarflexion torque encountered during experimental validation testing). Unlike hip testing, a low-level PD control was utilized during this test to capture the responsiveness of the system based on the closed-loop control scheme utilized for the ankle exoskeleton controller. The P and D gains were set to the same values as those determined via experimental tuning (P: 28, D: 200). Like the hip, five step response commands were performed while recording the duration, prescribed magnitude, and the measured torque of each step. Once the average prescribed magnitude and measured torque were determined across all five step responses, the rise time and overshoot/undershoot response of the system were calculated as described previously.

#### Torque Tracking

The accuracy of the prescribed ankle controller was characterized by evaluating the average root mean squared error between the prescribed and measured torques from two minutes of walking with closed loop ankle plantarflexion assistance (magnitude: 28 Nm).

#### Duration Testing

The operation length of the ankle configuration was examined while a user walked on a fixed-speed treadmill (1.25 ms^-1^) with maximum comfortable bilateral ankle plantarflexion assistance (28 Nm). As the user walked, the motor temperatures and battery voltage (from the 22.2v, 1800 mAh Li-Ion battery) were monitored on a minute-by-minute basis. The test lasted until one of the previously described criteria was met (motor temperatures or battery voltage reaching manufacturer’s recommended limits).

### Hip-&-Ankle Configuration

#### Duration Testing

Like the hip and ankle configurations individually, the length of operation of the combined hip-and-ankle configuration was tested while a user walked on a fixed-speed treadmill (1.25 ms^-1^) while receiving maximum comfortable joint assistance at both the hip and ankle joints (Hip Flexion: 5 Nm, Hip Extension: 5.5 Nm, Ankle Plantarflexion: 28 Nm). As they walked the motor temperatures for both joints of both sides and the battery voltage were recorded on a minute-by-minute basis until one of the previously described conditions for trial termination were met.

### Elbow Configuration

Complete information on the methodology used to validate the elbow configuration can be found in Colley et al. (*36*). The sections below provide a brief description of the methodologies utilized.

#### Benchtop Testing

Like the hip and ankle configurations, we tested the responsiveness of the elbow configuration via benchtop step-response testing. This consisted of locking the forearm and upper arm carbon fiber uprights to a table via vices. The same step response controller described previously was then applied with the maximum torque amplitude set to 10 Nm. A low-level PID control was utilized during this test to capture the responsiveness of the system based on the closed-loop control scheme utilized for the elbow exoskeleton controller. The P, I, and D gains were set to the same values as those determined via experimental tuning (P: 7, I: 25, D: 47). Eight step response commands were performed while recording the duration, prescribed magnitude, and the measured torque of each step. Once the average prescribed magnitude and measured torque were determined across all eight step responses, the rise time and overshoot/undershoot response of the system were calculated as described previously.

#### Torque Tracking

The accuracy of the prescribed elbow controller was characterized by evaluating the average root mean squared error between the prescribed and measured torques from thirteen consecutive 10 kg box lifting motions during closed-loop elbow assistance (magnitude: 12 Nm).

### Experimental Validation

To demonstrate the versatility and utility of our open-source system, we recruited healthy adults (**Supplemental Table 1**, n = 7) to complete activities while the device operated in different configurations. Inclusion criteria included age between 10-65, no lower-extremity orthopedic surgery within the prior six months, and no other health condition that would prevent safe completion of the protocol. Prior to enrollment all participants granted written informed consent of an Institutional Review Board approved protocol.

### Hip Configuration

To test the utility of the device while configured for hip assistance, we had two individuals (**Supplemental Table 1**; P1 & P2) walk on a treadmill inclined to 7.5 degrees. Prior to testing, both participants performed 60 minutes of hip assisted walking to acclimate to the device. This consisted of 30 minutes of level treadmill walking at a self-selected speed, and 30 minutes of incline treadmill walking (7.5°) at a self-selected speed that was subsequently maintained for testing. During acclimation, assistance in the extension and flexion directions were tuned based on participant comfort. Acclimation walks occurred the day prior to actual testing. On the test day, participants performed incline treadmill walking while outfitted with a portable, indirect calorimetry metabolic unit (K5, COSMED). Testing order was as follows: five minutes of quiet standing for a metabolic baseline, eight minutes of incline treadmill walking without the device (“shod”), fifteen minutes of rest, five minutes of quiet standing for a metabolic baseline, eight minutes of incline treadmill walking with hip assistance. Oxygen and carbon dioxide volumes during the last two minutes of treadmill incline walking were used to calculate steady-state metabolic power using Brockway’s equation (*40*). The respiratory data from quiet standing prior to each trial was used to calculate a baseline metabolic rate. The metabolic cost of transport for each trial was then calculated by offsetting the steady-state walking data by the baseline metabolic rate and normalizing it by body mass and walking speed.

### Ankle Configuration

One individual (**Supplemental Table 1**; P3), with prior ankle exoskeleton usage experience, completed ankle-assisted walking on outdoor terrain. This involved walking with, and without, the ankle exoskeleton about a relatively flat 1650 m loop at a local park (**Figure 6A&B**). Prior to testing, the individual completed one lap of the loop without exoskeleton assistance to familiarize them with the route. Ankle exoskeleton assistance was set to 30% of the user’s bodyweight (28 Nm). The testing order was as follows: one lap with exoskeleton assistance, twenty minutes of rest, one lap without wearing the exoskeleton device (“shod”). This order was selected as the device had mechanical failures during preliminary pilot testing and we wanted to avoid having the user waste time if a repeated failure were to occur during the actual test (it did not). The time to complete the loop and the number of steps taken with the right leg were recorded during testing. From these, we calculated the average walking speed of the user (dividing the length of the course by the time it took to complete it) and the average step length of the user (dividing the length of the course by the number of steps taken by the user). Metabolic data (oxygen and carbon dioxide volumes) over the course of each loop were collected; however, the data were collected in a way that made direct comparisons between conditions challenging (lack of distance markers for standardization, failure to record for a few minutes after completing the loop to account for the delay in metabolic response) and so this data was excluded as an outcome variable. This metabolic data has been made available for sake of transparency.

In addition to completing outdoor testing, this same participant completed metabolic testing with and without the exoskeleton device during level indoor treadmill walking. Prior to testing, the participant performed 60 minutes of ankle assisted treadmill walking to ensure acclimation to the device. This consisted of three, 20-minute sessions walking on the treadmill at 1.25 ms^-1^. Acclimation walks occurred the day prior to actual testing. On the test day, the participant performed level treadmill walking while outfitted with a portable, indirect calorimetry metabolic unit (K5, COSMED). Testing order was as follows: five minutes of quiet standing for a metabolic baseline, eight minutes of level treadmill walking without the device (“shod”), fifteen minutes of rest, five minutes of quiet standing for a metabolic baseline, eight minutes of level treadmill walking with ankle plantarflexion assistance. Oxygen and carbon dioxide volumes during the last two minutes of treadmill walking were used to calculate steady-state metabolic power using Brockway’s equation (*40*). The respiratory data from quiet standing prior to each trial was used to calculate a baseline metabolic rate. The metabolic cost of transport for each trial was then calculated by offsetting the steady-state walking data by the baseline metabolic rate and normalizing it by body mass and walking speed.

### Hip-&-Ankle Configuration

To test the utility of the device while configured for simultaneous hip-and-ankle assistance, we had two individuals (**Supplemental Table 1**; P4 & P5) walk on a treadmill while carrying a moderate sized load (22.5 %BW, weighted vest). Prior to testing, both participants performed 60 minutes of hip-and-ankle assisted walking to acclimate to the device. This consisted of 30 minutes of level treadmill walking without additional load carriage and 30 minutes of treadmill walking with the additional load. During acclimation, assistance in the extension and flexion directions of the hip and in the plantarflexion direction of the ankle were tuned based on participant comfort. Acclimation walks occurred the day prior to actual testing. Walking speed was held at 1.25 ms^-1^ on both the acclimation and testing visits. On the test day, participants performed load-carriage treadmill walking while outfitted with a portable, indirect calorimetry metabolic unit (K5, COSMED). Testing order was as follows: five minutes of quiet standing for a metabolic baseline, eight minutes of load-carriage treadmill walking with hip-and-ankle assistance, fifteen minutes rest, five minutes of standing metabolic baseline, eight minutes of load-carriage treadmill walking without the device (“shod”). Oxygen and carbon dioxide volumes during the last two minutes of treadmill walking were used to calculate steady-state metabolic power using Brockway’s equation (*40*). The respiratory data from quiet standing prior to each trial was used to calculate a baseline metabolic rate. The metabolic cost of transport for each trial was then calculated by offsetting the steady-state walking data by the baseline metabolic rate and normalizing it by body mass and walking speed.

### Elbow Configuration

To test the utility of the device while configured for elbow flexion assistance, we had two individuals (**Supplemental Table 1**; P6 & P7) perform weight curls with a 19.5 kg object until they reached fatigue and could not continue. The object was a wooden box with additional weights fixed to the inside (**Figure 7A**). A metronome set to sixty beats per minute was used to prompt users to raise and lower the weight (leading to a full cycle every two seconds). A repetition was considered one complete cycle from the down position back to the down position. Testing order was randomized across both participants resulting in the following test orders: P6 - no exoskeleton, twenty-five minutes of rest, exoskeleton and P7 - exoskeleton, twenty-five minutes of rest, no exoskeleton. The number of repetitions completed for each condition was recorded as the primary outcome of interest. Prior to testing, surface EMG electrodes were placed over the muscle belly of the short head of the biceps brachii muscle of each user’s dominate arm. Before performing fatigue lifting, both users performed a maximum voluntary isometric contraction (MVIC) by maximally flexing their forearm while holding a static strap with the elbow at 90^◦^. This value was then used to normalize the EMG data collected during fatigue lifting. The EMG data was analyzed by applying a 4^th^ order Butterworth band-pass filter (20-460 Hz), rectifying the signal, and then low-pass filtering the data with a 12 Hz cutoff. The EMG signal was normalized to each lifting cycle by identifying the troughs in the signal at the beginning and end of each cycle and then averaging across all the cycles for each condition (Exo, No Exo). Full information on the experimental testing protocol can be found in (*36*).

### Data Analysis

Given the small sample sizes for each experimental validation configuration, no statistical analyses were performed. All variables of interest were compared between the shod and exoskeleton conditions by calculating percent change, defined as:

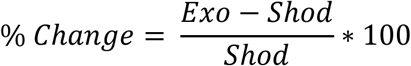

## Acknowledgments

We thank Dr. Karl Harshe, Dr. Pamela Bosch, and Dr. Collin Bowersock for their assistance with some of the experimental data collections.

## Author contributions

Conceptualization: JRW, CFC, PP, ZFL

Methodology: JRW, CFC, SF, DC, NE, PC, PP, ZFL

Investigation: JRW, CFC, SF, DC, NE, PC, PP, ZFL

Visualization: JRW, NE, PC, ZFL

Funding acquisition: ZFL

Project administration: JRW, ZFL

Supervision: JRW, ZFL

Writing – original draft: JRW

Writing – review & editing: JRW, CFC, SF, DC, NE, PC, PP, ZFL

## Data and materials availability

All data, code, and materials used in the analysis are available for other researchers to use, replicate, and analyze. Additionally, all material described above (software, electronics, hardware) and their documentation for OpenExo (in their state at the time of publication) are included with this manuscript to further facilitate easy user access. The data and version of the OpenExo at the time of manuscript submission can be found at: <u>DOI:</u> 10.5281/zenodo.13760782. Up-to-date versions of this material can be found on OpenExo’s website (theopenexo.org).

